# T-cell Abca1 and Abcg1 cholesterol efflux pathways suppress T-cell apoptosis and senescence and increase atherosclerosis in middle-aged *Ldlr*^-/-^ mice

**DOI:** 10.1101/2022.04.10.487770

**Authors:** Venetia Bazioti, Anouk M. La Rose, Sjors Maassen, Frans Bianchi, Rinse de Boer, Emma Guilbaud, Arthur Flohr-Svendsen, Anouk G. Groenen, Alejandro Marmolejo-Garza, Mirjam H. Koster, Niels J. Kloosterhuis, Alle T. Pranger, Miriam Langelaar-Makkinje, Alain de Bruin, Bart van de Sluis, Alison B. Kohan, Laurent Yvan-Charvet, Geert van den Bogaart, Marit Westerterp

**Affiliations:** Department of Pediatrics, University Medical Center Groningen, University of Groningen, Groningen, 9713 AV, the Netherlands; Department of Molecular Immunology and Microbiology, Groningen Biomolecular Sciences and Biotechnology Institute, University of Groningen, Groningen, 9747 AG, the Netherlands; Institut National de la Santé et de la Recherche Médicale (INSERM) U1065, Université Côte d’Azur, Centre Méditerranéen de Médecine Moléculaire (C3M), Atip-Avenir, Fédération Hospitalo-Universitaire (FHU) Oncoage, 06204 Nice, France; European Research Institute for the Biology of Ageing, University Medical Center Groningen, University of Groningen, Groningen, 9713 AV, the Netherlands; Laboratory of Medicine, University Medical Center Groningen, University of Groningen, Groningen, 9713 AV, the Netherlands; Department of Biomolecular Health Sciences, Dutch Molecular Pathology Center, Faculty of Veterinary Medicine, Utrecht University, Utrecht, 3584 CL, the Netherlands; Division of Endocrinology and Metabolism, Department of Medicine, University of Pittsburgh, Pittsburgh, PA 15260, United States

## Abstract

Atherosclerosis is a chronic inflammatory disease driven by hypercholesterolemia. During aging, T-cells accumulate cholesterol, which could lead to a pro-inflammatory phenotype. However, the role of cholesterol efflux pathways mediated by ATP-binding cassette A1 and G1 (ABCA1/ABCG1) in T-cell-dependent age-related inflammation and atherosclerosis remains poorly understood. In this study, we generated mice with T-cell-specific *Abca1/Abcg1*-deficiency on the low-density-lipoprotein-receptor deficient (*Ldlr*^-/-^) background. T-cell *Abca1/Abcg1*-deficiency decreased blood, lymph node, and splenic T-cells, and increased T-cell activation and apoptosis. T-cell *Abca1/Abcg1*-deficiency induced a premature T-cell aging phenotype in middle-aged (12-13 months) *Ldlr*^-/-^ mice, reflected by upregulation of senescence markers. Despite T-cell senescence and enhanced T-cell activation, T-cell *Abca1/Abcg1*-deficiency decreased atherosclerosis and aortic inflammation in middle-aged *Ldlr*^-/-^ mice, accompanied by decreased T-cells in atherosclerotic plaques. We attribute these effects to T-cell apoptosis downstream of T-cell activation. Collectively, T-cell cholesterol efflux pathways are critical for maintaining T-cell numbers, suppress senescence, and induce atherosclerosis in middle-aged *Ldlr*^-/-^ mice.

## INTRODUCTION

Atherosclerosis is a lipid-driven chronic inflammatory disease of the large- and mid-sized arteries that can lead to myocardial infarction or stroke ^1^. T-cells play an important role in atherosclerosis, by secreting pro- and anti-inflammatory cytokines that affect plaque formation ^2,3^. While initially most T-cells in atherosclerotic plaques were reported to be of the T helper 1 (T_h_1) phenotype that is pro-atherogenic due to secretion of the cytokines interferon gamma (IFNγ) and tumor necrosis factor alpha (TNFα) ^4–6^, later studies have shown that regulatory T-cells (T_regs_) can be anti-atherogenic by secreting transforming growth factor beta (TGFβ) and interleukin 10 (IL-10) ^7–9^. More recent studies revealed that during atherosclerosis and cardiovascular disease (CVD), T_regs_ acquire pro-inflammatory characteristics of T_h_1 and T follicular helper (T_FH_) cells ^10–14^. Feeding a cholesterol-rich Western-type diet (WTD) to mice deficient in apolipoprotein E (*Apoe^-/-^*) induces a phenotypic switch from T_regs_ to T_h_1 and T_FH_ cells ^11^. This phenotypic switch was prevented by injections of apolipoprotein A-I (apoA-I), which stimulates cholesterol efflux. Hence, these data suggest a key role for T-cell cholesterol accumulation in T_regs_ acquiring pro-inflammatory characteristics ^11^. Moreover, increased plasma membrane cholesterol accumulation in CD8^+^ T-cells deficient in the enzyme Acetyl-CoA cholesterol Acyltransferase 1 (ACAT1) that esterifies cholesterol, enhances differentiation of naïve T-cells into IFNγ producing T-cells ^15^, presumably with pro-atherogenic effects. Together, these findings suggest that exacerbated cholesterol accumulation in T-cells increases T-cell subsets with pro-inflammatory characteristics that enhance atherosclerosis, but this has not been studied directly.

T-cells from aged (~73 years old) compared to young (~22 years old) humans also show increased membrane cholesterol accumulation ^16^. During aging, the number of naïve T-cells decreases, while activated T-cells that show features of cellular senescence and secrete pro-inflammatory cytokines increase ^17–19^. This process may contribute to inflammaging ^20,21^, the pro-inflammatory state that develops during aging. Aging is a major risk factor for atherosclerosis, presumably due to inflammaging ^22,23^. Although thymic atrophy may contribute to the decrease in naïve T-cells and increase in activated T-cells with features of cellular senescence that promote inflammaging ^20,24^, data in mice with T-cell *Acat1* deficiency or *Apoe* deficiency ^11,15^ suggest that cholesterol accumulation in T-cells during aging may also directly promote secretion of pro-inflammatory cytokines.

The cholesterol transporters ATP Binding Cassette A1 and G1 (ABCA1 and ABCG1) mediate cholesterol efflux to apoA-I and high-density-lipoprotein (HDL), respectively ^25–27^. Previous studies have shown that activation of the T-cell receptor (TCR) by anti-CD3, which increases T-cell proliferation, decreases expression of *Abca1* and *Abcg1* by >90% ^28^, leading to T-cell plasma membrane cholesterol accumulation ^28,29^. It has been suggested that *Abcg1* is the most highly expressed cholesterol transporter in T-cells ^28^. T-cell *Abcg1* deficiency increases differentiation of naïve T-cells into T_regs_, which suppresses atherosclerosis ^30^. However, *Abcg1*^-/-^ T-cells show a 6-fold increase in *Abca1* expression ^31^, suggesting that similar to macrophages ^32^, the cholesterol transporters ABCA1 and ABCG1 have overlapping roles and show mutual compensation in T-cells. We here investigated whether combined deficiency of *Abca1* and *Abcg1* in T-cells would exacerbate the cholesterol accumulation phenotype, and thus impact T-cell subsets, T-cell aging, and atherogenesis.

## RESULTS

### T-cell *Abca1/Abcg1* Deficiency Increases Cholesterol Accumulation

We generated a T-cell specific *Abca1/Abcg1*-deficient mouse model by crossbreeding *Abca1^fl/fl^Abcg1^fl/fl^* mice with mice expressing the T-cell specific *CD4Cre* promoter. The *CD4Cre* promoter starts to be expressed at the double positive (DP) (CD4^+^CD8^+^) stage of thymic T-cell maturation and results in deletion of *loxP* flanked genes in the double positive (DP) and single positive (SP) (CD4^+^ or CD8^+^) T-cell populations ^33,34^. To assess the deletion of *Abca1* and *Abcg1* in T-cells, we isolated splenic T-cells, since the yield of thymic SP T-cells was too low to properly assess the expression of these transporters. In *CD4CreAbca1^fl/fl^Abcg1^fl/fl^* splenic T-cells, *Abca1* and *Abcg1* mRNA expression were decreased by >90% compared to *Abca1^fl/fl^Abcg1^fl/fl^* T-cells (Fig. 1a). To assess the potential role of T-cell ABCA1 and ABCG1 mediated cholesterol efflux pathways in atherosclerosis, *CD4CreAbca1^fl/fl^Abcg1^fl/fl^* mice were crossbred with *Ldlr*^-/-^ mice to generate *CD4CreAbca1^fl/fl^Abcg1^fl/fl^Ldlr*^-/-^ mice and *Abca1^fl/fl^Abcg1^fl/fl^Ldlr*^-/-^ controls. We refer to these mice as *T-Abc^dko^Ldlr*^-/-^ and *Ldlr*^-/-^ mice, respectively.

**Fig. 1.**
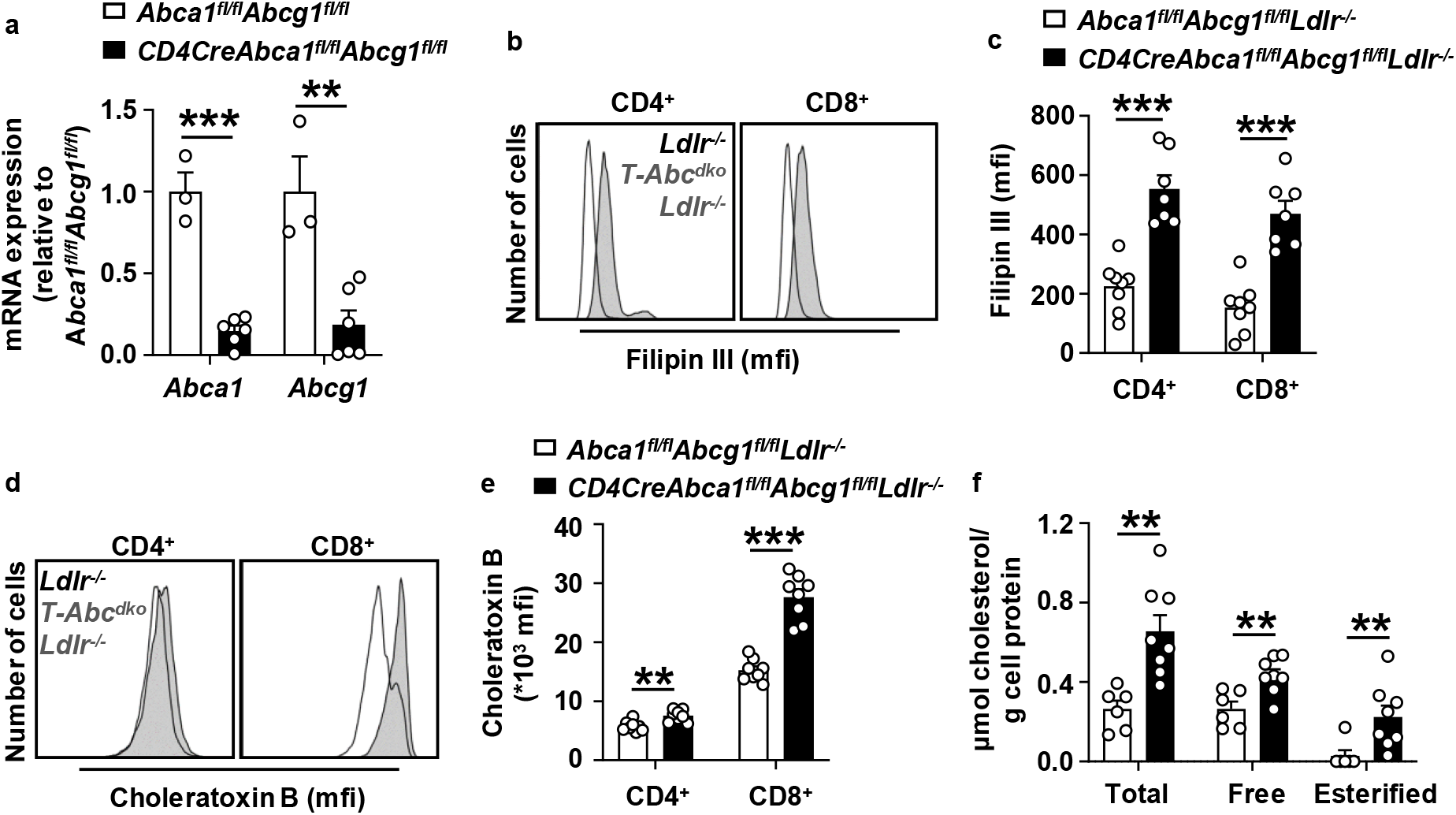
T-cell *Abca1/Abcg1* deficiency increases cholesterol accumulation. *CD4CreAbca1^fl/fl^Abcg1^fl/fl^*, *Abca1^fl/fl^Abcg1^fl/fl^*, *CD4CreAbca1^fl/fl^Abcg1^fl/fl^Ldlr*^-/-^, and *Abca1^fl/fl^Abcg1^fl/fl^Ldlr*^-/-^ mice were fed a chow diet. (**a**) Splenic T-cells were isolated from *CD4CreAbca1^fl/fl^Abcg1^fl/fl^* and *Abca1^fl/fl^Abcg1^fl/fl^* mice and RNA was extracted. *Abca1* and *Abcg1* mRNA expression were measured by qPCR. n=3-6. (**b-e**) Filipin and choleratoxin B stainings on blood CD4^+^ and CD8^+^ T-cells from *CD4CreAbca1^fl/fl^Abcg1^fl/fl^Ldlr*^-/-^ and *Abca1^fl/fl^Abcg1^fl/fl^Ldlr*^-/-^ mice were assessed by flow cytometry. (**b**) Representative flow cytometry plots of filipin staining and (**c**) quantification. n=7-8. (**d**) Representative flow cytometry plots of choleratoxin B staining and (**e**) quantification. n=8. (**f**) Splenic T-cells were isolated from *CD4CreAbca1^fl/fl^Abcg1^fl/fl^Ldlr*^-/-^ and *Abca1^fl/fl^Abcg1^fl/fl^Ldlr*^-/-^ mice, and total and free cholesterol were measured by GC-MS. Esterified cholesterol equals total cholesterol - free cholesterol. n=6-8. For all panels, error bars represent standard error of the mean (SEM). n indicates biological replicates. *p* value was determined by unpaired two-tailed Student’s t-test. \*\**p*<0.01, \*\*\**p*<0.001.

To assess the functional consequences of T-cell *Abca1/Abcg1* deficiency, we performed filipin staining to measure free cholesterol accumulation. T-cell *Abca1/Abcg1* deficiency increased filipin staining in thymic DP, CD4^+^, and CD8^+^ T-cells (Supplementary Fig. 1a, b), and in CD4^+^ and CD8^+^ T-cells in blood (Fig. 1b, c). We next assessed the cellular localization of cholesterol in splenic T-cells by confocal microscopy. T-cell *Abca1/Abcg1* deficiency increased filipin staining at the plasma membrane, reflecting plasma membrane cholesterol accumulation (Supplementary Fig. 1c). We also observed increased staining of choleratoxin B, suggestive of increased ganglioside GM1, a component of lipid rafts (Fig. 1d, e).

To validate the findings on cholesterol accumulation, we performed Gas Chromatography – Mass Spectrometry (GC-MS). T-cell *Abca1/Abcg1* deficiency increased total cholesterol by 2.5-fold, reflected by increases in free cholesterol and cholesteryl esters (CE) (Fig. 1f). In line with CE accumulation, T-cell *Abca1/Abcg1* deficiency increased Oil Red O staining, reflecting lipid droplets in para-aortic lymph nodes (LNs) (Supplementary Fig. 1d, e), and BODIPY 493/503 staining in splenic T-cells (Supplementary Fig. 1f). To assess whether cell organelles were affected in *Abca1/Abcg1*-deficient T-cells, we performed transmission electron microscopy. We confirmed the presence of large lipid droplets in *Abca1/Abcg1*-deficient CD4^+^ and CD8^+^ T-cells, and otherwise observed no overt differences (Supplementary Fig. 1g, h).

Collectively, deficiency of *Abca1* and *Abcg1* in T-cells increased free and esterified cholesterol, reflected by plasma membrane cholesterol accumulation and presence of lipid droplets.

### T-cell *Abca1/Abcg1* Deficiency Leads to Plasma Membrane Stiffening

High concentrations of plasma membrane cholesterol increase plasma membrane stiffness in model membranes and cells ^35,36^. To examine whether plasma membrane stiffness was affected by T-cell *Abca1/Abcg1* deficiency, we stained T-cells with the fluorescent dye BODIPY C10 and performed Fluorescence-Lifetime Imaging Microscopy (FLIM). BODIPY C10 is a molecular rotor that has stiffness-dependent fluorescence lifetime ^37^. When BODIPY C10 binds to a fluid membrane, it shows high rotational speed which decreases its fluorescence lifetime, while binding to a stiff membrane leads to increased fluorescence lifetime ^37^. *Abca1/Abcg1*-deficient CD4^+^ and CD8^+^ T-cells showed increased BODIPY C10 fluorescence lifetime, compared to control CD4^+^ and CD8^+^ T-cells (Fig. 2a, b and Supplementary Fig. 2a-d), suggesting that T-cell *Abca1/Abcg1* deficiency increased cell stiffness. In *Abca1/Abcg1*-deficient CD4^+^ and CD8^+^ T-cells structures that resemble CE-rich lipid droplets showed yellow staining which reflects very high fluorescence lifetime (Fig. 2a, b). When we excluded these from analysis, BODIPY C10 fluorescence lifetime was still increased in *Abca1/Abcg1*-deficient CD4^+^ and CD8^+^ T-cells compared to control CD4^+^ and CD8^+^ T-cells (Fig. 2c-f). These data suggest that accumulation of cholesterol in *Abca1/Abcg1-* deficient T-cells leads to plasma membrane stiffening.

**Fig. 2.**
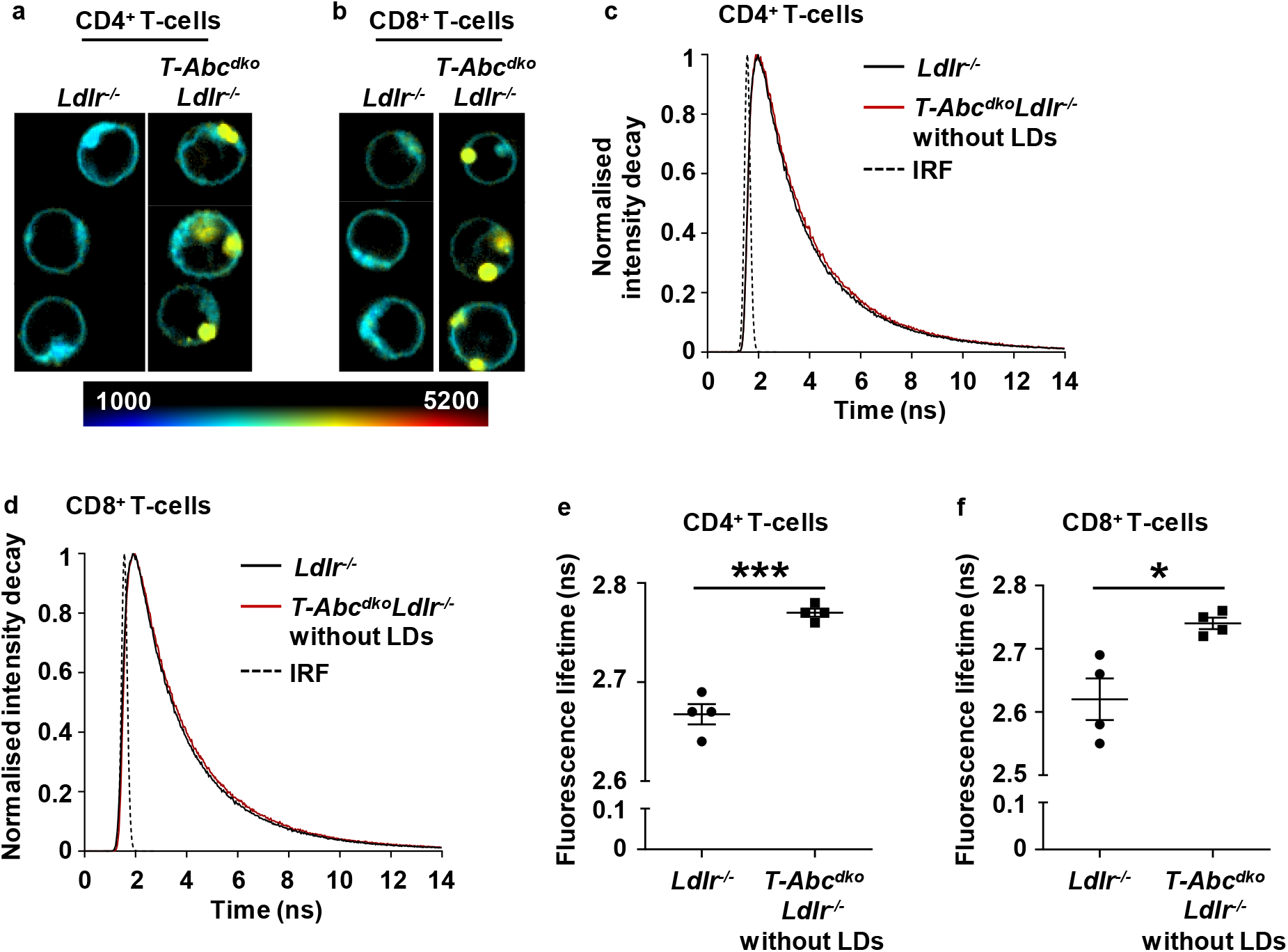
T-cell *Abca1/Abcg1* deficiency leads to plasma membrane stiffening. *T-Abc^dko^Ldlr*^-/-^ and *Ldlr*^-/-^ mice were fed a chow diet. Spleens were collected and CD4^+^ and CD8^+^ T-cells were isolated. CD4^+^ and CD8^+^ T-cells were stained with BODIPY C10 and analyzed by Fluorescence-Lifetime Imaging Microscopy (FLIM). Representative images for CD4^+^ (**a**) and CD8^+^ (**b**) T-cells are shown. Fluorescence lifetime decay curves for CD4^+^ (**c**) and CD8^+^ (**d**) T-cells are shown and quantified in the plasma membrane (without LDs) (**e-f**). Each data point represents an individual mouse. LD = lipid droplet. (**c-d**) Dashed lines: fits with mono-exponential decay functions convoluted with the instrument response function (IRF). n=4. For all panels, error bars represent SEM. n indicates biological replicates. *p* value was determined by unpaired two-tailed Student’s t-test. \**p*<0.05, \*\*\**p*<0.001.

### T-cell *Abca1/Abcg1* Deficiency Decreases Peripheral T-cells and Increases T-cell Activation

It has been shown previously that *LckCreAbca1^fl/fl^Abcg1^fl/fl^* mice with T-cell specific *Abca1/Abcg1* deficiency have decreased thymic CD4^+^ and CD8^+^ T-cells compared to littermate controls and decreased splenic CD4^+^ and CD8^+^ T-cells ^38^. We thus assessed whether T-cell *Abca1/Abcg1* deficiency affected the number of T-cells in *T-Abc^dko^Ldlr*^-/-^ mice. While single T-cell *Abca1* or *Abcg1* deficiency did not affect T-cell numbers (Supplementary Fig. 3a), combined T-cell *Abca1/Abcg1* deficiency decreased blood CD4^+^ and CD8^+^ T-cells by ~50% both in *Ldlr*^-/-^ (Fig. 3a, b) and *Ldlr^+/+^* mice (Supplementary Fig. 3b). Because of these decreases, we evaluated thymic T-cell populations. T-cell *Abca1/Abcg1* deficiency did not affect thymic weight (data not shown). *T-Abc^dko^Ldlr*^-/-^ mice showed no changes in thymic T-cell populations compared to littermate controls (Supplementary Fig. 3c-f). We next examined T-cell numbers in spleen and para-aortic LNs. Similar to observations in blood, we found that T-cell *Abca1/Abcg1* deficiency decreased splenic and para-aortic LN CD4^+^ and CD8^+^ T-cells by ~50-70% (Fig. 3c, d). Together these data indicate that T-cell *Abca1/Abcg1* deficiency decreases blood, splenic, and para-aortic LN T-cells without affecting thymic T-cells.

**Fig. 3.**
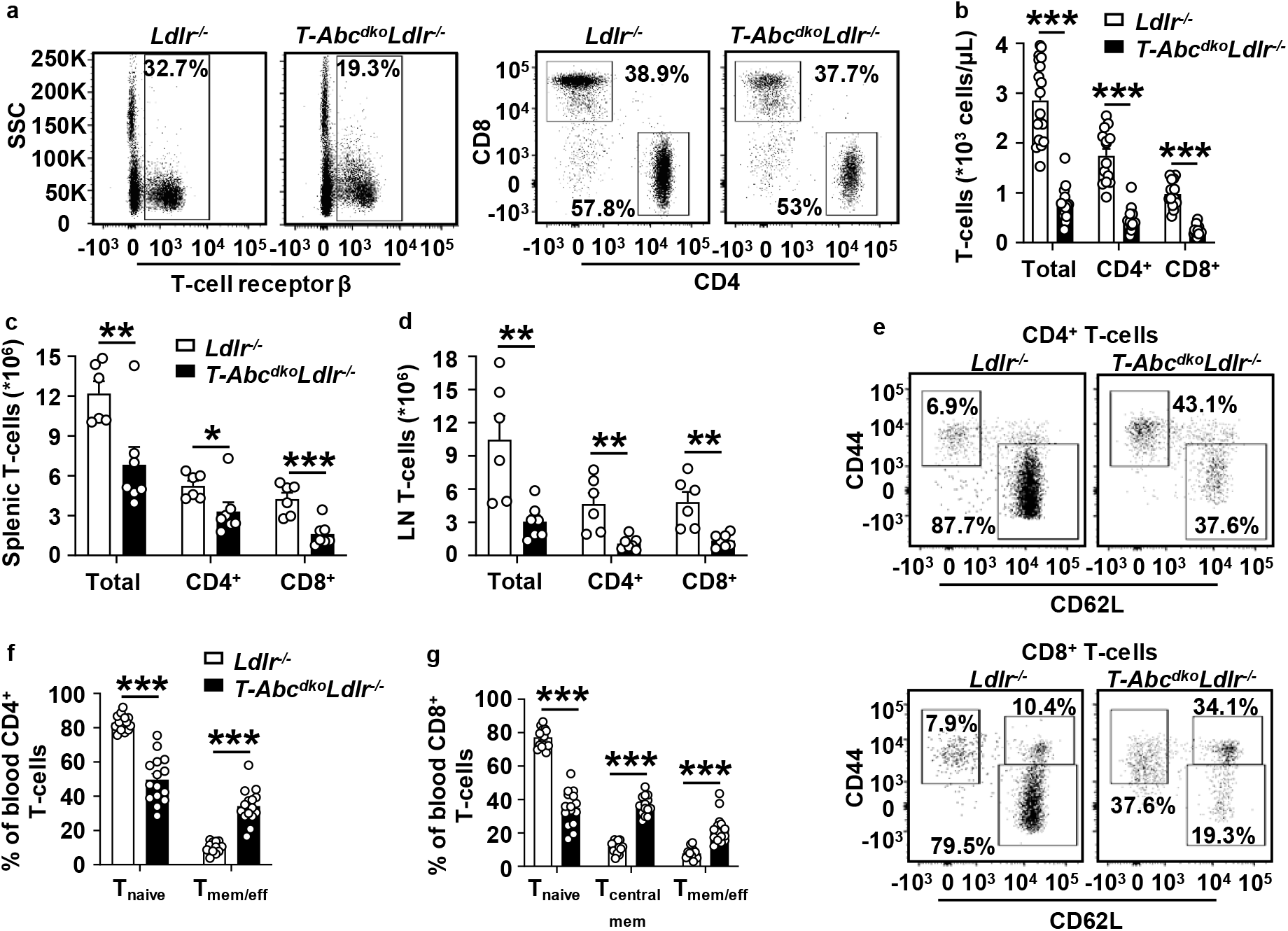
T-cell *Abca1/Abcg1* deficiency decreases blood, splenic, and para-aortic lymph node T-cells and increases T-cell activation. *Ldlr*^-/-^ and *T-Abc^dko^Ldlr*^-/-^ mice were fed a chow diet. Blood, spleens, and para-aortic lymph node (LNs) were collected. Cells were stained with the indicated antibodies and analyzed by flow cytometry. (**a-b**) Gating strategy (**a**) and quantification (**b**) of TCRβ^+^ (total), CD4^+^, and CD8^+^ T-cells in blood. n=15-18. (**c-d**) Total, CD4^+^, and CD8^+^ T-cells in spleen (**c**) and para-aortic LNs (**d**). n=6-7. (**e-g**) Representative flow cytometry dot plots (**e**) and quantification of CD4^+^ CD44^-^CD62L^+^ (T_naive_) and CD44^+^CD62L^-^ (T_mem/eff_) (**f**), and CD8^+^ T_naive_, T_mem/eff_, and CD8^+^CD44^+^CD62L^+^ (T_central mem_) (**g**) cells in blood. n=14-15. For all panels, error bars represent SEM. n indicates biological replicates. *p* value was determined by unpaired two-tailed Student’s t-test. \**p*<0.05, \*\**p*<0.01, \*\*\**p*<0.001.

We have previously shown that T-cell *Abca1/Abcg1* deficiency mildly enhanced CD4^+^ T-cell activation in the *LckCreAbca1^fl/fl^Abcg1^fl/fl^* model ^39^. We thus assessed the effect of single and combined T-cell *Abca1/Abcg1* deficiency on T-cell activation. Single T-cell *Abca1* or *Abcg1* deficiency did not affect T-cell activation in blood (Supplementary Fig. 3g, h). Combined T-cell *Abca1/Abcg1* deficiency increased T_memory/effector_ (CD4^+^CD44^+^CD62L^-^ and CD8^+^CD44^+^CD62L^-^) and T_central memory_ (CD8^+^CD44^+^CD62L^+^) cells by >3-fold (Fig. 3e-g). Further, T-cell *Abca1/Abcg1* deficiency decreased CD4^+^ and CD8^+^ T_naive_ cells (CD44^-^CD62L^+^) cells by ~50% (Fig. 3e-g). We observed similar effects in spleen and para-aortic LNs (Supplementary Fig. 3i-l). Collectively, as a percentage of total CD4^+^ or CD8^+^ T-cells, T-cell *Abca1/Abcg1* deficiency increased T_memory/effector_ and T_central memory_ cells in blood, spleen, and para-aortic LNs. However, when corrected for total number of T-cells, the increases in T_memory/effector_ and T_central memory_ cells were no longer significant between the genotypes (Supplementary Fig. 3m-q), and CD4^+^ T_memory/effector_ cells were decreased in para-aortic LNs (Supplementary Fig. 3q). Moreover, T-cell *Abca1/Abcg1* deficiency decreased CD4^+^ and CD8^+^ T_naive_ cells by >70% (Supplementary Fig. 3m-p, r). In sum, while increasing the percentage of CD4^+^ and CD8^+^ T-cells in the memory/effector and central memory subsets, T-cell *Abca1/Abcg1* deficiency dramatically decreased T_naive_ cells in blood, spleen, and para-aortic LNs.

### Effects of T-cell *Abca1/Abcg1* Deficiency on T-cell Exhaustion

Previous studies in T-cells have shown that T-cell cholesterol accumulation induces T-cell exhaustion, which was mainly attributed to cholesterol accumulation in the ER ^40^. Exhausted T-cells are prone to apoptosis and show reduced proliferative capacity ^40,41^. We therefore investigated whether increased cholesterol accumulation due to loss of *Abca1/Abcg1* in T-cells would induce T-cell exhaustion and thus explain the decrease in T-cell numbers.

The percentage of para-aortic LN CD4^+^PD-1^+^ T-cells was increased in *T-Abc^dko^Ldlr*^-/-^ mice compared to *Ldlr*^-/-^ controls (Supplementary Fig. 4a), while not being affected after correction for total T-cell numbers (Supplementary Fig. 4b). T-cell *Abca1/Abcg1* deficiency did not affect the expression of the exhaustion markers TIM3 and LAG3 and induced only a modest increase in CTLA4 on CD4^+^ and CD8^+^ T-cells (Supplementary Fig. 4c-h). T-cell *Abca1/Abcg1* deficiency increased the transcription factor Eomes in CD8^+^ T-cells, an exhaustion marker (Supplementary Fig. 4i, j) ^42^. However, increased Eomes has also been associated with an increase in T_central memory_ cells ^43^. The increase in PD-1 on CD4^+^ T-cells may be due to increased T-cell activation ^44,45^. We observed similar effects on splenic CD4^+^PD-1^+^ T-cell numbers and expression of CTLA4 and Eomes in splenic T-cells (data not shown). Since CTLA4 was minimally increased by T-cell *Abca1/Abcg1* deficiency, and the increase in PD-1 and Eomes could be attributed to the expansion of other T-cell populations, these data are suggestive of only minor effect of T-cell *Abca1/Abcg1* deficiency on T-cell exhaustion.

### T-cell *Abca1/Abcg1* Deficiency Increases T-cell Apoptosis Upon TCR Stimulation

We then investigated other mechanisms contributing to the decrease in peripheral T-cell numbers. Previous studies have shown that stimulation of the TCR by anti-CD3 decreases the expression of *Abca1* and *Abcg1* by >90%, while upregulating the expression of *Acat1*, 3-hydroxy-3-methyl-glutaryl-coenzyme A reductase (*Hmgcr*), and *Ldlr* ^28^. These effects were confirmed by a later study ^15^, and suggest that TCR stimulation induces a change in gene expression that favors cholesterol accumulation, likely to generate substrates for cellular growth and proliferation. We confirmed that TCR stimulation decreased *Abca1* and *Abcg1* mRNA expression in T-cells (Supplementary Fig. 5a, b). In genetic models of plasma membrane cholesterol accumulation in T-cells, TCR signaling is enhanced ^15,28,31^. Conversely, TCR signaling is suppressed in T-cells deficient in SCAP (Sterol regulatory Element Binding Protein (SREBP) cleavage activating protein) that cannot synthesize cholesterol ^46^. We thus assessed whether T-cell *Abca1/Abcg1* deficiency affected T-cell proliferation downstream of TCR signaling. Upon TCR stimulation, *Abca1/Abcg1* deficiency increased T-cell proliferation in CD4^+^ and CD8^+^ T-cells (Supplementary Fig. 5c-e). Even though in line with observations of increased T-cell proliferation in T-cell *Abcg1* deficiency ^28,31^, these findings cannot explain the decreased peripheral T-cells in *T-Abc^dko^Ldlr^-/-^* mice compared to *Ldlr*^-/-^ controls. However, these data strongly suggest that T-cell *Abca1/Abcg1* deficiency enhances TCR signaling. We thus investigated whether T-cell *Abca1/Abcg1* deficiency affects other pathways that regulate peripheral T-cell numbers, downstream of TCR signaling.

TCR stimulation enhances the differentiation of T_naive_ cells into T_memory/effector_ and T_central memory_ cells ^47^. The increase in T_memory/effector_ and T_central memory_ cells in mice with T-cell *Abca1/Abcg1* deficiency (Fig. 3e-g and Supplementary Fig. 3i-l) is consistent with increased TCR signaling. Upon TCR stimulation, T-cells differentiate into T_memory/effector_ cells that may undergo apoptosis, in a pathway known as activation-induced cell death (AICD) ^48^. Using anti-CD3 and IL-2 as stimuli, we examined whether T-cell *Abca1/Abcg1* deficiency enhanced apoptosis downstream of TCR stimulation, by monitoring expression of cleaved Caspase3/7 in T-cells over time using the Incucyte system. T-cell *Abca1/Abcg1* deficiency increased Caspase3/7^+^ expression by 3.7-fold in CD4^+^ T-cells and 1.8-fold in CD8^+^ T-cells, reflecting increased T-cell apoptosis (Fig. 4a, b and Supplementary Fig. 5f).

**Fig. 4.**
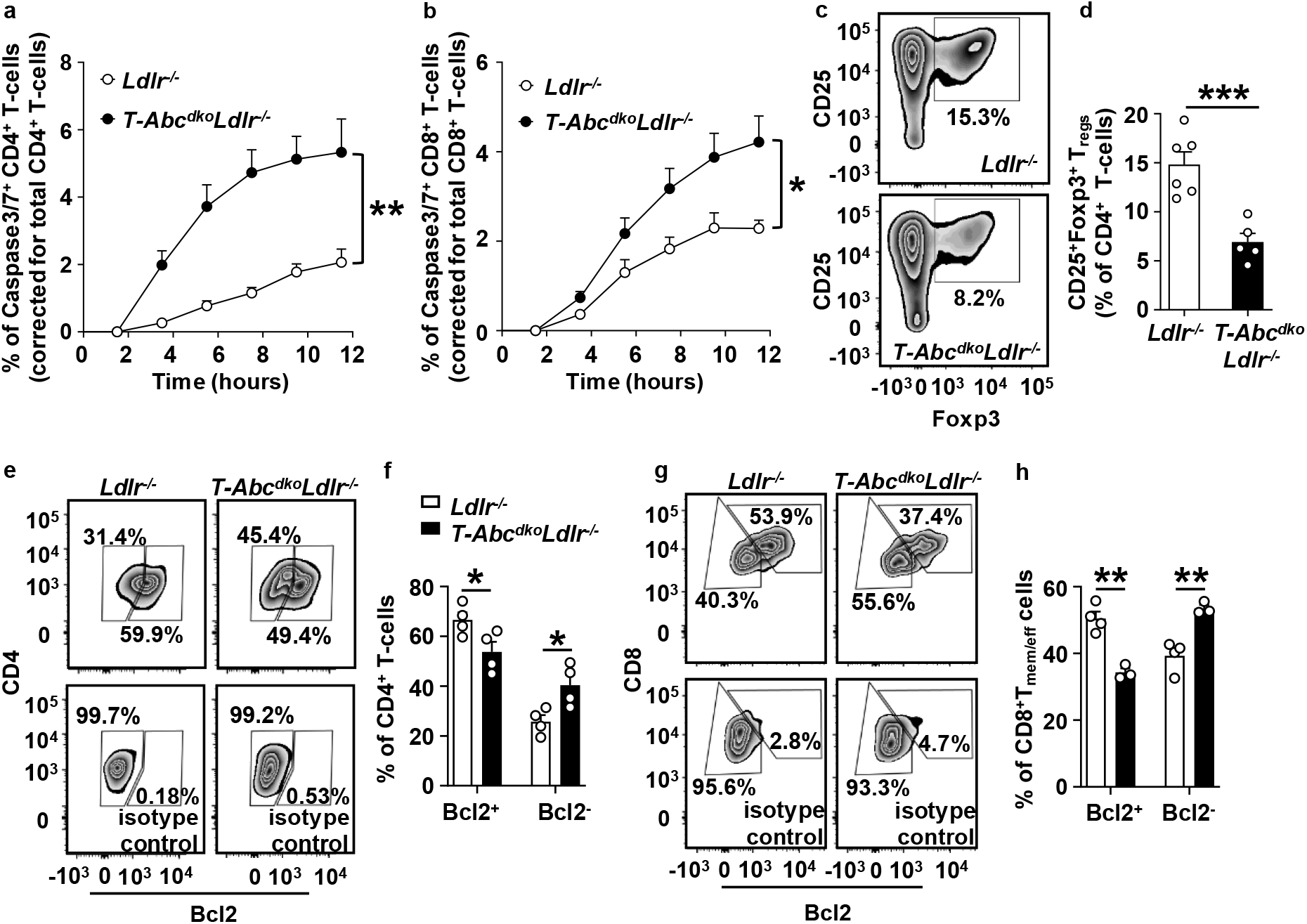
T-cell *Abca1/Abcg1* deficiency increases T-cell apoptosis. *T-Abc^dko^Ldlr*^-/-^ and *Ldlr^-/-^* mice were fed a chow diet. Spleens were collected and CD4^+^ and CD8^+^ T-cells were isolated. (**a-b**) CD4^+^ and CD8^+^ T-cells were stimulated with αCD3 and IL-2 for 12 h with concomitant staining for cleaved Caspase3/7. CD4^+^ (**a**) and CD8^+^ (**b**) T-cells acquiring Caspase 3^+^/7^+^ staining over time were assessed using the Incucyte system. n=7-8. (**c-d**) CD4^+^ T-cells were stimulated with αCD3/αCD28 beads, IL-2, and TGFβ for 72 h. Representative flow cytometry plots of CD25^+^Foxp3^+^ T_regs_ are shown (**c**), corrected for their respective isotype control, and quantified (**d**). n=5-6. (**e-h**) CD4^+^ and CD8^+^ T-cells were stimulated with αCD3 and IL-2 for 12 h and stained for Bcl2 (**e-f**) and CD44 and CD62L (**g-h**) after stimulation. (**e-f**) Representative flow cytometry plots of CD4^+^Bcl2^+^ and CD4^+^Bcl2^-^ T-cells are shown (**e**), corrected for their respective isotype control, and quantified (**f**). n=4. (**g-h**) Representative flow cytometry plots of CD8^+^ T_memory/effector_ Bcl2^+^ and Bcl2^-^ cells are shown (**g**), corrected for their respective isotype control, and quantified (**h**). n=3-4. For all panels, error bars represent SEM. n indicates biological replicates. *p* value was determined by unpaired two-tailed Student’s t-test. \**p*<0.05, \*\**p*<0.01, \*\*\**p*<0.001.

Given that *Abcg1* deficiency alone enhances differentiation of T_naive_ cells in T_regs_ ^30^, and T_regs_ have an athero-protective role, we then also studied T_reg_ differentiation in T-cells with combined *Abca1/Abcg1* deficiency. We used TGFβ, anti-CD3, and IL-2 to induce CD25^+^Foxp3^+^ T_reg_ differentiation, similar to previous studies ^30,49^, and found that, in contrast to T-cell *Abcg1* deficiency ^30^, T-cell *Abca1/Abcg1* deficiency suppressed T_reg_ differentiation by ~50% (Fig. 4c, d). Given that, with the exception of TGFβ, stimuli for the apoptosis assay were similar to those used for T_reg_ differentiation, it is likely that the decrease in T_regs_ is simply the consequence of increased T-cell apoptosis. We thus studied mechanisms that cause T-cell apoptosis.

We found that the expression of the death receptor FAS that is highly expressed in lipid rafts and involved in the extrinsic apoptosis pathway ^50^ was not affected by T-cell *Abca1/Abcg1* deficiency upon stimulation with anti-CD3 and IL-2 (Supplementary Fig. 5g, h). We next measured the expression of the anti-apoptotic protein Bcl2 that regulates the intrinsic apoptosis pathway ^51^. Upon anti-CD3/IL-2 stimulation, T-cell *Abca1/Abcg1* deficiency decreased anti-apoptotic Bcl2^+^CD4^+^ T-cells by ~20%, while increasing pro-apoptotic Bcl2^-^CD4^+^ T-cells by ~60% (Fig. 4e, f). T-cell *Abca1/Abcg1* deficiency did not affect Bcl2 expression in all CD8^+^ T-cells (data not shown), likely because of the expanded long-lived CD8^+^ T_central memory_ cell population ^52^. Since AICD implies apoptosis of the T_memory/effector_ cell population, we studied the effect of T-cell *Abca1/Abcg1* deficiency on Bcl2 expression in T_memory/effector_ cells. Upon anti-CD3/IL-2 stimulation, T-cell *Abca1/Abcg1* deficiency decreased anti-apoptotic Bcl2^+^CD8^+^ T_memory/effector_ cells by ~30%, while increasing pro-apoptotic Bcl2^-^CD8^+^ T_memory/effector_ cells by ~40% (Fig. 4g, h). Once the Bcl2 pathway is activated, mitochondrial ROS (mitoROS) stimulates cytochrome c release from mitochondria, which ultimately leads to caspase activation ^53^. Indeed, upon anti-CD3/IL-2 stimulation, T-cell *Abca1/Abcg1* deficiency increased mitoROS levels, mainly in CD8^+^ T_naive_ and T_memory/effector_ cells (Supplementary Fig. 5i-l). There was no effect on CD4^+^ T-cells (Supplementary Fig. 5i, j), perhaps because T-cells with high mitoROS had already undergone apoptosis. Collectively, these data indicate that T-cell *Abca1/Abcg1* deficiency enhances T-cell apoptosis downstream of TCR and CD25, mediated by Bcl2 in the intrinsic apoptosis pathway, especially in T_memory/effector_ cells. This likely accounts for the ~50% decrease in peripheral T-cells in T-cell *Abca1/Abcg1* deficiency.

### Aging Increases Cholesterol Accumulation and Apoptosis in T-cells from Wild-Type Mice

We then asked whether there would be a broader physiological relevance of the phenotype we observed in T-cell *Abca1/Abcg1* deficiency. Aged humans (~73 years old) show increased T-cell plasma membrane cholesterol accumulation compared to young humans (~22 years old) ^16^ and a decrease in peripheral T-cells ^17–19^. The phenotype in T-cells from aged individuals resembles the phenotype of mice with T-cell *Abca1/Abcg1* deficiency and may suggest a broader physiological relevance for T-cell plasma membrane cholesterol accumulation in regulating peripheral T-cell numbers. Using T-cells from aged (~24 months old) and young (~3 months old) mice, we investigated this further.

Similar to findings in aged humans ^16^, we found that blood CD4^+^ and CD8^+^ T-cells from aged mice show an increase in filipin staining compared to young mice, reflecting an increase in plasma membrane cholesterol accumulation (Fig. 5a, b). In line with previous observations from aged humans and mice ^17,18,54^, aged mice showed a decrease in peripheral T-cells compared to young mice (Fig. 5c), mainly due to decreased T_naive_ cells, while T_memory/effector_ and T_central memory_ cells were increased (Fig. 5d, e). CD4^+^ T-cells from aged mice showed a slight decrease in T-cell proliferation in response to TCR stimulation, while CD8^+^ T-cell proliferation was not affected by aging (Fig. 5f-h), in line with previous findings showing that T_naive_ cells from aged mice still proliferate efficiently ^18^. We then studied apoptosis in response to TCR stimulation in young and aged T-cells. TCR stimulation increased cleaved Caspase3/7 in aged compared to young T-cells (Fig. 5i, j), reflecting increased apoptosis. The data in aged mice (Fig. 5a-e, i, j) together with our findings in *T-Abc^dko^Ldlr*^-/-^ mice suggest that plasma membrane cholesterol accumulation in T-cells from aged mice increases T-cell activation and apoptosis, which likely contributes to the decrease in T-cells during aging.

**Fig. 5.**
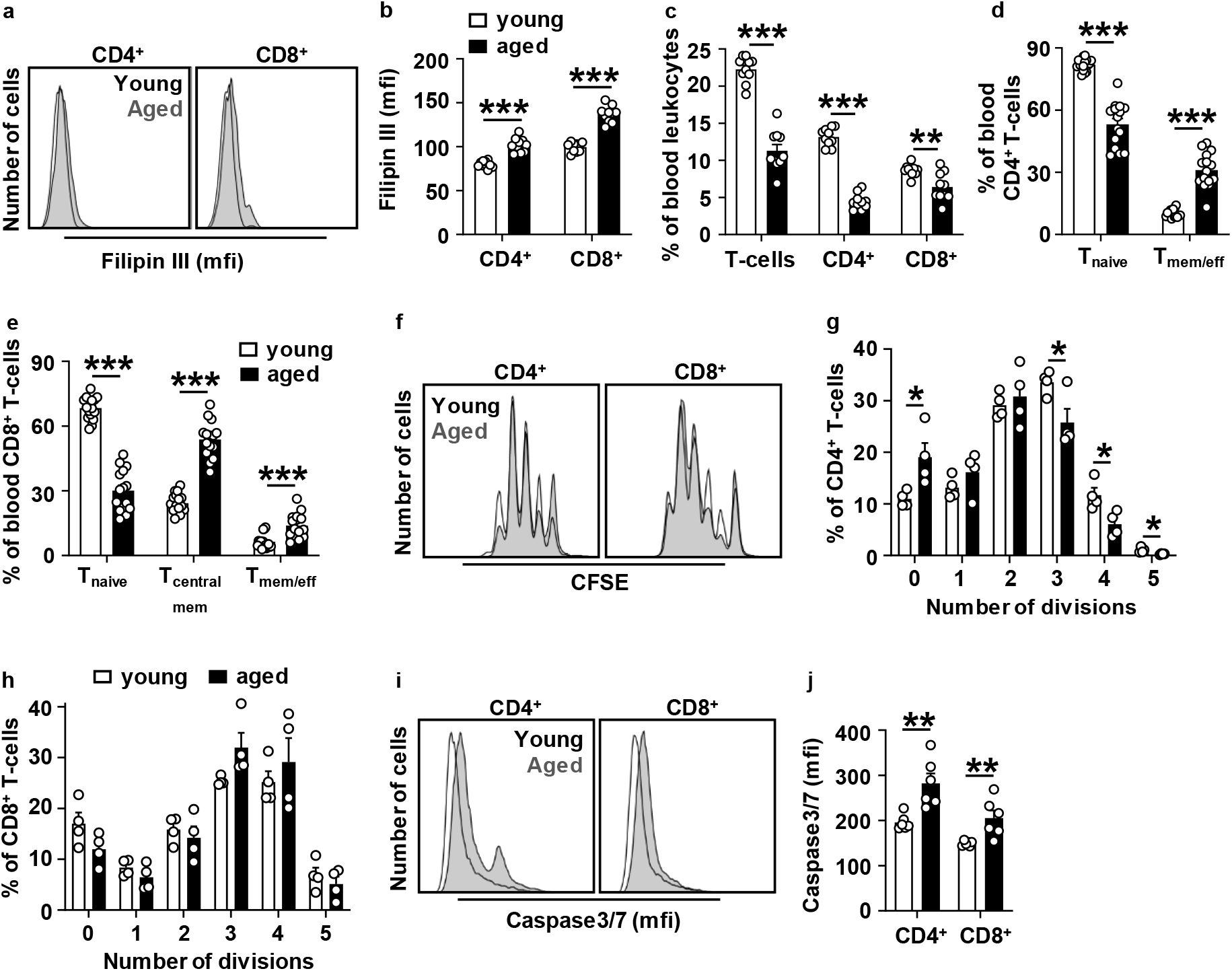
Aging increases cholesterol accumulation and apoptosis in T-cells from wild-type mice. Blood and spleens from young (3 months) and aged (24 months) wild-type mice fed a chow diet were collected. (**a-b**) Filipin staining on blood CD4^+^ and CD8^+^ T-cells was assessed by flow cytometry. (**a**) Representative flow cytometry plots and (**b**) quantification. n=10. (**c**) Total, CD4^+^, and CD8^+^ T-cells in blood. n=10. (**d-e**) CD4^+^ T_naive_ and T_mem/eff_ cells (**d**) and CD8^+^ T_naive_, T_mem/eff_, and T_central mem_ cells (**e**) in blood. n=15-16. (**f-j**) Splenic CD4^+^ and CD8^+^ T-cells were isolated. (**f-h**) T-cells were labeled with CFSE and stimulated with αCD3/αCD28 beads. CFSE dilution was measured at 72 h after stimulation by flow cytometry. Representative CFSE dilutions are shown (**f**) and the number of divisions were quantified for CD4^+^ (**g**) and CD8^+^ (**h**) T-cells. n=4. (**i-j**) CD4^+^ and CD8^+^ T-cells were stimulated with αCD3 and IL-2 for 12 h with concomitant staining for cleaved Caspase3/7. Representative flow cytometry plots of cleaved Caspase3/7 expression in CD4^+^ and CD8^+^ T-cells are shown (**i**) and quantified (**j**). n=6. For all panels, error bars represent SEM. n indicates biological replicates. *p* value was determined by unpaired two-tailed Student’s t-test. \**p*<0.05, \*\**p*<0.01, \*\*\**p*<0.001.

### T-cell *Abca1/Abcg1* Deficiency Induces a Premature T-Cell Aging Phenotype

We then asked whether aging would affect TCR responses in *T-Abc^dko^Ldlr*^-/-^ mice. We aged *T-Abc^dko^Ldlr*^-/-^ mice and *Ldlr*^-/-^ littermate controls until 12-13 months (middle-aged). The phenotype in terms of T-cell numbers and activation did not differ between middle-aged and young *T-Abc^dko^Ldlr*^-/-^ mice (data not shown). T-cell *Abca1/Abcg1* deficiency increased cleaved Caspase3/7 upon TCR stimulation in middle-aged mice, reflecting increased apoptosis (Fig. 6a, b). Strikingly, upon TCR stimulation, T-cell *Abca1/Abcg1* deficiency almost completely abolished CD4^+^ and CD8^+^ T-cell proliferation, while T-cells from control mice still proliferated (Fig. 6c-e). T-cells with *Abca1/Abcg1* deficiency may not enter the cell cycle due to upregulation of cyclin dependent kinase inhibitors, including p21 ^55^. *Abca1/Abcg1* deficiency increased *p21* mRNA expression by 6-fold in CD4^+^ T-cells and 2.5-fold in CD8^+^ T-cells compared to middle-aged control T-cells (Fig. 6f, g), while *p16^INK4A^* mRNA expression was not affected (data not shown). Of note, unlike in other cell types, in T-cells, *p16^INK4A^* rather reflects increased T-cell activation than aberrant cell cycling ^19^. When comparing *p21* mRNA expression in T-cells from middle-aged *T-Abc^dko^Ldlr*^-/-^ mice to T-cells from aged wild-type mice, we observed that *Abca1/Abcg1* deficiency increased *p21* mRNA expression by ~6-fold in CD4^+^ and CD8^+^ T-cells (Fig. 6f, g). Together, these data indicate that middle-aged *Abca1/Abcg1*-deficient T-cells upregulate *p21* and do not enter the cell cycle, which is suggestive of T-cell senescence and premature T-cell aging.

**Fig. 6.**
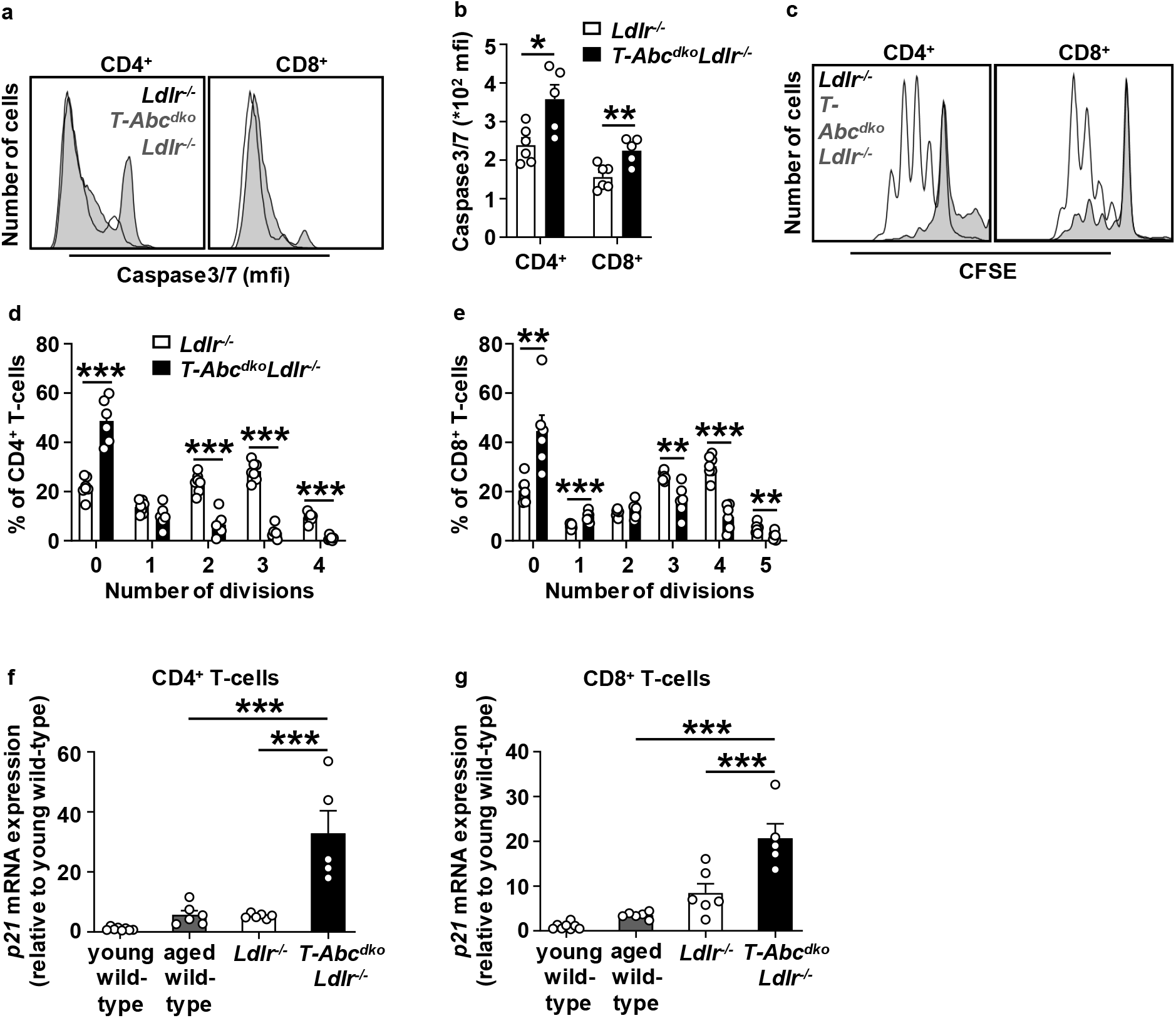
T-cell *Abca1/Abcg1* deficiency induces a premature T-cell aging phenotype. *T-Abc^dko^Ldlr*^-/-^ and *Ldlr*^-/-^ mice were fed a chow diet for 12-13 months (**a-g**) and wild-type mice for 3 months (young) or 24 months (aged) (**f-g**). (**a-g**) Spleens were collected and CD4^+^ and CD8^+^ T-cells were isolated. (**a-b**) CD4^+^ and CD8^+^ T-cells from *T-Abc^dko^Ldlr*^-/-^ and *Ldlr*^-/-^ mice were stimulated with αCD3 and IL-2 for 12 h with concomitant staining for cleaved Caspase3/7. Representative flow cytometry plots of cleaved Caspase3/7 expression in CD4^+^ and CD8^+^ T-cells are shown (**a**) and quantified (**b**). n=5-6. (**c-e**) T-cells from *T-Abc^dko^Ldlr*^-/-^ and *Ldlr*^-/-^ mice were labeled with CFSE and stimulated with αCD3/αCD28 beads. CFSE dilution was measured at 72 h after stimulation by flow cytometry. Representative CFSE dilutions are shown (**c**) and the number of divisions were quantified for CD4^+^ (**d**) and CD8^+^ (**e**) T-cells. n=6-7. (**f-g**) T-cells from *T-Abc^dko^Ldlr^-/-^, Ldlr^-/-^*, and wild-type mice were isolated and RNA was extracted. *p21* mRNA expression was measured in CD4^+^ (**f**) and CD8^+^ (**g**) T-cells by qPCR and expressed as fold change compared to young wild-type mice. n=5-8. For all panels, error bars represent SEM. n indicates biological replicates. *p* value was determined by unpaired two-tailed Student’s t-test (**b, d-e**) or one-way ANOVA with Bonferroni post-test (**f-g**). \**p*<0.05, \*\**p*<0.01, \*\*\**p*<0.001.

### Effects of T-cell *Abca1/Abcg1* Deficiency on T-cell Subsets and Atherosclerosis

Previous studies have shown that T-cell *Abcg1* deficiency decreases atherosclerosis by enhancing the formation of T_regs_ in thymus and LNs of *Ldlr*^-/-^ mice fed WTD ^30^. In addition, WTD feeding enhances the conversion of T_regs_ into T_FH_ and T_h_1 cells, which was dependent on cholesterol accumulation ^11^. T-cell *Acat1* deficiency induces plasma membrane cholesterol accumulation in CD8^+^ T-cells, leading to expansion of CD8^+^IFNγ^+^ T-cells ^11,15^ that have a pro-atherogenic role. We thus investigated whether exacerbated cholesterol accumulation due to loss of *Abca1/Abcg1* in T-cells would affect T-cell subsets.

We assessed these T-cell subsets in para-aortic LNs of *T-Abc^dko^Ldlr*^-/-^ and *Ldlr*^-/-^ mice fed a chow diet. While the percentage of para-aortic LN CD25^+^Foxp3^+^ T_regs_ was not affected, T-cell *Abca1/Abcg1* deficiency increased the percentage of CD25^-^Foxp3^+^ T_regs_ (Fig. 7a), which may suggest that upon cholesterol accumulation, T_regs_ lose their CD25 expression, and start to express markers of T_FH_ and T_h_1 cells as has been reported in *Apoe^-/-^* mice ^11^. Both in *T-Abc^dko^Ldlr*^-/-^ and *Ldlr*^-/-^ mice, CD25^-^Foxp3^+^ T-cells were mainly of the activated T_memory/effector_ (CD44^+^CD62L^-^) phenotype, while CD25^+^Foxp3^+^ cells were mostly of the T_naive_ (CD44^-^CD62L^+^) phenotype (Supplementary Fig. 6a, b). After correction for total T-cell numbers, T-cell *Abca1/Abcg1* deficiency decreased CD25^+^Foxp3^+^ T_regs_ by ~75%, while not affecting CD25^-^Foxp3^+^ T_regs_ (Fig. 7b), consistent with the decrease in CD25^+^Foxp3^+^ T_regs_ that we observed *in vitro* (Fig. 4c, d), presumably due to increased T-cell apoptosis. T-cell *Abca1/Abcg1* deficiency increased the percentage of T_FH_ cells in para-aortic LNs (Fig. 7c), but did not affect T_FH_ cells after correction for total T-cell numbers (Fig. 7d).

**Fig. 7.**
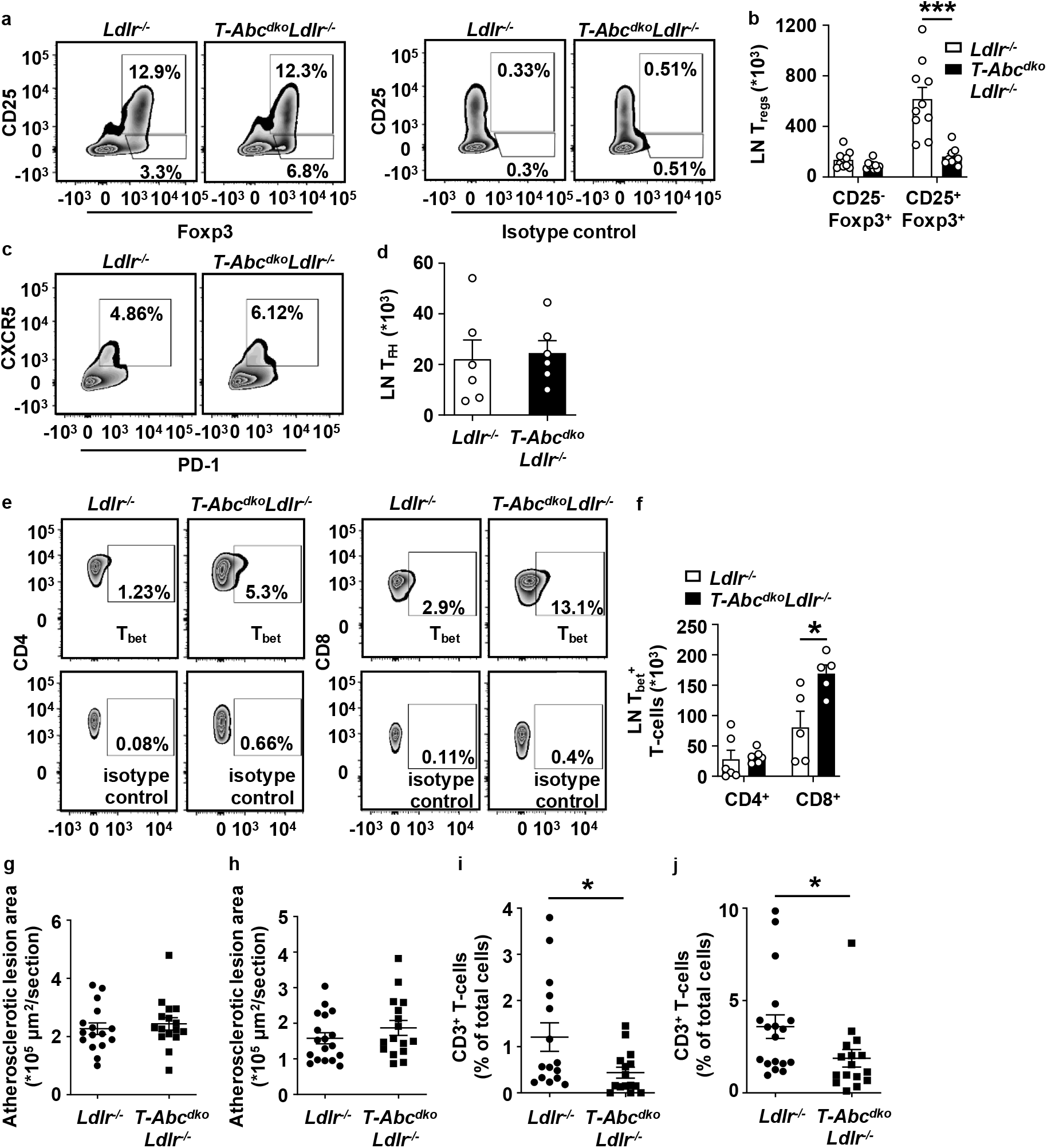
Effects of T-cell *Abca1/Abcg1* deficiency on T-cell subsets in para-aortic LNs and atherosclerosis in *Ldlr-/-* mice. Female *T-Abc^dko^Ldlr*^-/-^ and *Ldlr*^-/-^ mice were fed a chow diet for 28 weeks (**a-f, h, j**) or a Western-type diet (WTD) for 10 weeks (**g, i**). (**a-f**) Para-aortic LNs were isolated and cells were stained with the indicated antibodies and analyzed by flow cytometry. (**a-b**) Representative flow cytometry plots of CD25^-^Foxp3^+^ and CD25^+^Foxp3^+^ T_regs_ are shown (**a**), corrected for their respective isotype control and total number of LN cells, and quantified (**b**). n=9-10. (**c-d**) Representative flow cytometry plots of CD4^+^CD44^+^CD62L^-^CXCR5^+^PD1^+^ T_FH_ cells are shown (**c**), corrected for total number of LN cells, and quantified (**d**). n=6-7. (**e-f**) Representative flow cytometry plots of CD4^+^ and CD8^+^T_bet_^+^ T-cells are shown (**e**), corrected for their respective isotype control and total number of LN cells, and quantified (**f**). n=5-6. (**g-j**) Hearts were isolated and sections of the aortic root were prepared and stained for haematoxylin-eosin (H&E) (**g-h**) or CD3 (**i-j**). Atherosclerotic lesion area of WTD-fed (**g**) and chow diet-fed (**h**) mice. CD3^+^ cells per section were quantified and corrected for total number of cells for WTD-fed (**i**) and chow diet-fed (**j**) mice. n=14-18. Each data point represents an individual mouse. For all panels, error bars represent SEM. n indicates biological replicates. *p* value was determined by unpaired two-tailed Student’s t-test. \**p*<0.05, \*\*\**p*<0.001.

Plasma membrane cholesterol accumulation increases formation of T_h_1 cells that express the transcription factor T_bet_ ^11,15^. The percentage of CD4^+^ and CD8^+^ T_bet_^+^ cells was increased in para-aortic LNs of *T-Abc^dko^Ldlr*^-/-^ mice compared to *Ldlr*^-/-^ controls (Fig. 7e) and not affected after correction for total T-cell numbers for CD4^+^T_bet_^+^ cells, while CD8^+^T_bet_^+^ cells were still ~2-fold increased (Fig. 7f). While CD4^+^T_bet_^+^ cells were mainly of the T_memory/effector_ (CD44^+^CD62L^-^) phenotype (Supplementary Fig. 6c), CD8^+^T_bet_^+^ cells were mostly of the long-lived T_central memory_ (CD44^+^CD62L^+^) phenotype (Supplementary Fig. 6d). In sum, T-cell *Abca1/Abcg1* deficiency decreased CD25^+^Foxp3^+^ T_regs_ and increased CD8^+^ T_bet_^+^ cells in para-aortic LNs, while para-aortic LN CD25^-^Foxp3^+^, T_FH_, and T_h_1 cells were not affected (Table 1).

**Table 1.** Summary of effects of T-cell *Abca1/Abcg1* deficiency on T-cell subsets in para-aortic LNs from *Ldlr*^-/-^ mice fed chow diet.

| T-cell subset | Cell numbers |
| --- | --- |
| CD25<sup>+</sup>Foxp3<sup>+</sup> T<sub>regs</sub><br>(~55% T<sub>naive</sub>) | ~75% ↓ |
| CD25<sup>-</sup>Foxp3<sup>+</sup> T<sub>regs</sub><br>(~60% T<sub>memory/effector</sub>) | ↔ |
| T<sub>FH</sub> | ↔ |
| CD4<sup>+</sup>T<sub>bet</sub><sup>+</sup><br>(~55% T<sub>memory/effector</sub>) | ↔ |
| CD8<sup>+</sup>T<sub>bet</sub><sup>+</sup><br>(~75% T<sub>central memory</sub>) | ~2-fold ↑ |

Para-aortic LNs were isolated and cells were stained with the indicated antibodies and analyzed by flow cytometry. Data are shown in Figure 7.

We then investigated the effects of these changes on atherosclerosis. To induce atherogenesis, female *T-Abc^dko^Ldlr*^-/-^ mice and *Ldlr*^-/-^ littermate controls were fed a WTD. Findings on T-cell activation and T-cell subsets were similar to mice fed chow diet (data not shown). T-cell *Abca1/Abcg1* deficiency did not affect blood monocytes or neutrophils (Supplementary Fig. 6e). After 10 weeks of WTD, atherosclerotic lesion size was assessed at the level of the aortic root. T-cell *Abca1/Abcg1* deficiency did not affect atherosclerotic lesion size (Fig. 7g and Supplementary Fig. 6f). This was accompanied by a decrease in plasma total cholesterol levels of ~15% (Supplementary Table 1), which reflects decreased very low-density-lipoprotein (VLDL) and low-density-lipoprotein (LDL) cholesterol (Supplementary Fig. 6g). In an attempt to exclude the confounding factor of decreased VLDL/LDL cholesterol levels to atherogenesis, we fed mice a chow diet for 28 weeks, similar to a previous study ^56^. Chow diet-fed *T-Abc^dko^Ldlr*^-/-^ and *Ldlr*^-/-^ mice had similar plasma total cholesterol levels (Supplementary Table 1) and developed atherosclerotic lesions similar in size compared to WTD-fed *Ldlr*^-/-^ mice (Supplementary Fig. 6h), but there was no difference in atherosclerotic lesion size between the genotypes (Fig. 7h). In line with observations in blood and secondary lymphoid organs, *T-Abc^dko^Ldlr*^-/-^ mice showed a ~50% decrease in plaque CD3^+^ T-cells on both diets compared to *Ldlr*^-/-^ controls (Fig. 7i, j and Supplementary Fig. 6i, j).

Hence, despite decreasing T-cells in atherosclerotic plaques and decreasing para-aortic LN CD25^+^Foxp3^+^ T_regs_, and increasing para-aortic LN CD8^+^ T_bet_^+^ cells (Table 1), T-cell *Abca1/Abcg1* deficiency did not affect atherosclerotic lesion size in *Ldlr*^-/-^ mice fed chow diet or WTD. Since these subsets have both pro- and anti-inflammatory effects, and the loss of *Abca1/Abcg1* in T-cells does not result in a single predominant subset, this effect on atherosclerosis could be due to counter-regulatory inflammatory effects in the aorta during sterile inflammation.

### Effects of T-cell *Abca1/Abcg1* Deficiency on T-cell Subsets and Atherosclerosis in Middle-Aged *Ldlr-/-* Mice

We then investigated whether the premature T-cell aging phenotype in middle-aged *T-Abc^dko^Ldlr*^-/-^ mice affected atherogenesis. T-cell *Abca1/Abcg1* deficiency in middle-aged *Ldlr*^-/-^ mice did not affect blood monocytes or neutrophils (Supplementary Fig. 7a). Similar to observations in young *T-Abc^dko^Ldlr*^-/-^ mice (Fig. 7a-f), CD25^+^Foxp3^+^ T_regs_ were decreased by ~70% (Fig. 8a and Supplementary Fig. 7b) and CD4^+^ T_memory/effector_ by ~50% (Fig. 8b), while CD8^+^T_bet_^+^ cells were increased by ~2.5-fold (Fig. 8c and Supplementary Fig. 7c) in para-aortic LNs from middle-aged *T-Abc^dko^Ldlr*^-/-^ mice compared to controls. CD25^-^Foxp3^+^ T_regs_ (Fig. 8a and Supplementary Fig. 7b), CD4^+^T_bet_^+^ cells (Fig. 8c and Supplementary Fig. 7c), and T_FH_ cells (Fig. 8d and Supplementary Fig. 7d) did not differ between the genotypes. Under conditions of similar plasma total cholesterol levels (Supplementary Table 1), T-cell *Abca1/Abcg1* deficiency decreased atherosclerotic lesion size by ~35% (Fig. 8e, f), which was accompanied by a ~50% decrease in plaque CD3^+^ T-cells (Fig. 8g, h). While the extent of the decrease in plaque CD3^+^ T-cells was similar to our observations in young mice (Fig. 7i, j), the decrease in atherosclerotic lesion area was not, and also the atherosclerotic lesions of middle-aged mice showed a 2-fold higher T-cell content than those of young mice (Fig. 7j), perhaps suggesting a more prominent role of T-cells in atherosclerosis in middle-aged *Ldlr*^-/-^ mice.

**Fig. 8.**
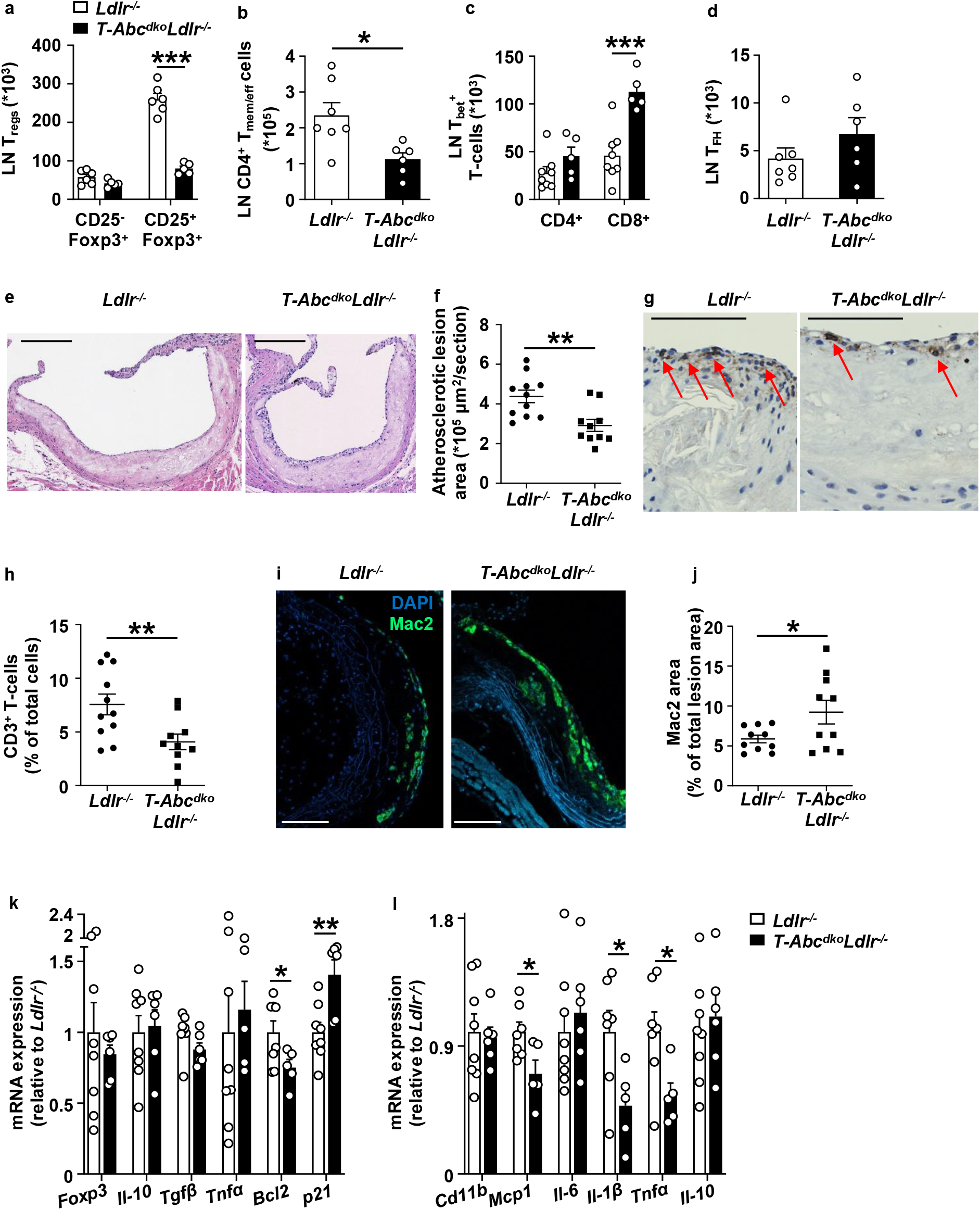
T-cell *Abca1/Abcg1* deficiency decreases atherosclerosis in middle-aged *Ldlr*^-/-^ mice. *T-Abc^dko^Ldlr*^-/-^ and *Ldlr*^-/-^ mice were fed a chow diet for 12-13 months. **(a-d)** Para-aortic LNs were isolated and cells were stained with the indicated antibodies and analyzed by flow cytometry. (**a**) Numbers of CD25^-^Foxp3^+^ and CD25^+^Foxp3^+^ T_regs_ corrected for their respective isotype control. n=5-6. (**b**) Numbers of CD4^+^ T_mem/eff_ T-cells. n=6-7. (**c**) Numbers of CD4^+^ and CD8^+^T_bet_^+^ T-cells corrected for their respective isotype control. n=5-8. (**d**) Numbers of T_FH_ cells. n=6-7. (**e-i**) Hearts were isolated and sections of the aortic root were prepared and stained for H&E (**e-f**) or CD3 (**g-h**) or Mac2 and DAPI (**i-j**). (**e**) Representative pictures of H&E staining on the aortic root. Scale bar represents 200 μm. (**f**) Atherosclerotic lesion area. (**g**) Representative pictures of CD3 staining on the atherosclerotic lesion. T-cells were identified as cells with brown plasma membrane CD3 staining and the hematoxylin staining still visible. CD3^+^ cells are depicted by arrows. Scale bar represents 80 μm. (**h**) CD3^+^ cells per section were quantified and corrected for total number of cells. (**i**) Representative pictures of Mac2 staining on the aortic root. Scale bar represents 100 μm. (**j**) Mac2 area corrected for total atherosclerotic lesion area. n=8-11. (**f, h, j**) Each data point represents an individual mouse. (**k-l**) CD3^+^ T-cells and CD3^-^ cells were isolated from aortas, and RNA was extracted. (**k**) *Foxp3, Il-10, Tgfβ, Tnfα, Bcl2*, and *p21* mRNA expression in aortic CD3^+^ T-cells and (**l**) *Cd11b, Mcp1, Il-6, Il-1β, Tnfα*, and *Il-10* mRNA expression in aortic CD3^-^ cells. n=5-8. For all panels, error bars represent SEM. n indicates biological replicates. *p* value was determined by unpaired two-tailed Student’s t-test. \**p*<0.05, \*\**p*<0.01, \*\*\**p*<0.001.

To obtain more insights into the processes that affected atherogenesis in middle-aged *T-Abc^dko^Ldlr*^-/-^ mice, we characterized their atherosclerotic lesions further. Although the density of macrophages in atherosclerotic lesions was relatively low (~6-10% of the atherosclerotic lesions), T-cell *Abca1/Abcg1* deficiency did increase macrophage content by ~60% (Fig. 8i, j). Only few necrotic cores were present in atherosclerotic plaques and the necrotic core area was not different between the genotypes (Supplementary Fig. 7e). The relative increase in macrophages suggests less advanced atherosclerosis in middle-aged *T-Abc^dko^Ldlr*^-/-^ mice compared to their controls.

To further investigate mechanisms underlying the decrease in atherosclerosis in middle-aged *T-Abc^dko^Ldlr*^-/-^ mice, we isolated T-cells from the whole aorta. Unlike our findings on the decrease of CD25^+^Foxp3^+^ T_regs_ in para-aortic LNs, T-cell *Abca1/Abcg1* deficiency did not affect mRNA expression of the T_reg_ transcription factor *Foxp3* (Fig. 8k). Also, mRNA expression of anti-inflammatory *Il-10* and *Tgfβ*, and pro-inflammatory *Tnfα* in aortic T-cells did not differ between the genotypes (Fig. 8k). In line with findings in splenic T-cells (Fig. 6f, g), T-cell *Abca1/Abcg1* deficiency increased mRNA expression of the senescence marker *p21* in aortic T-cells, and decreased *Bcl2* mRNA by ~25% (Fig. 8k). The decrease in Bcl2 suggests that also in the aorta, T-cells from middle-aged *T-Abc^dko^Ldlr*^-/-^ mice are prone to apoptosis. We then investigated whether T-cell *Abca1/Abcg1* deficiency would affect inflammatory gene expression in other cell types of the aorta. As such, we characterized the T-cell negative fraction of the aorta, consisting of endothelial cells, myeloid cells, and smooth muscle cells. While T-cell *Abca1/Abcg1* deficiency did not affect mRNA expression of the myeloid cell marker *Cd11b*, or the cytokines *Il-6* and *Il-10*, it decreased mRNA expression of monocyte chemoattractant protein 1 (*Mcp1*), *Il-1β*, and *Tnfα*, which have pro-atherogenic effects (Fig. 8l). It has been reported that uptake of apoptotic T-cells by macrophages may suppress pro-inflammatory gene expression ^57–60^. Hence, we attribute the decrease in atherosclerosis to T-cell apoptosis and uptake of apoptotic T-cells by myeloid cells, which suppresses aortic inflammation.

## DISCUSSION

These studies provide a link between defective cholesterol efflux and plasma membrane cholesterol accumulation and membrane stiffening in promoting T-cell activation and T-cell apoptosis with athero-protective effects in middle-aged mice (Fig. 9). Previous studies have shown that T-cells accumulate cholesterol during aging ^16^, and that T-cell aging induces T-cell activation and decreases peripheral T-cell numbers. Mechanistically, we now found that T-cell *Abca1/Abcg1* deficiency induced premature T-cell aging, reflected by an almost complete suppression of T-cell proliferation and expression of senescence markers (Fig. 9).

**Fig. 9.**
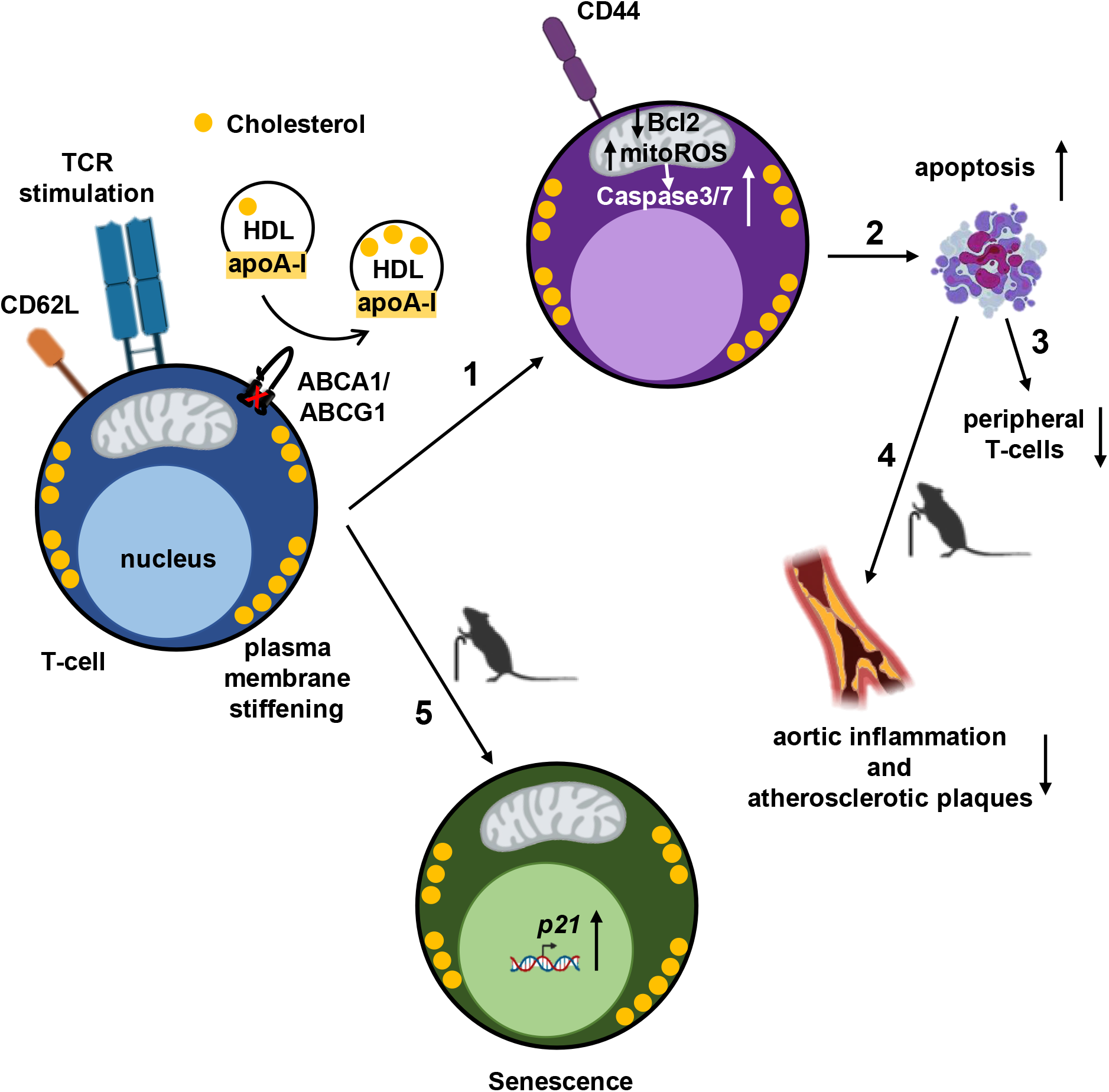
T-cell *Abca1/Abcg1* deficiency induces T-cell apoptosis and T-cell senescence in middle-aged mice. 1) T-cell *Abca1/Abcg1* deficiency increases T-cell activation, leading to 2) apoptosis, and 3) a decrease in peripheral T-cells, and 4) a decrease in aortic inflammation and atherosclerosis in middle-aged mice. 5) T-cell *Abca1/Abcg1* deficiency induces cellular senescence in middle-aged mice. TCR = T-cell receptor.

Association studies in the multi-ethnic study of atherosclerosis (MESA) have shown that a low level of CD4^+^ naïve T-cells in blood was associated with increased carotid intima media thickness (cIMT), suggesting that T-cell activation enhances atherogenesis in humans ^61^. T-cells are numerous in advanced atherosclerotic plaques of human carotids (~50-65% of all plaque cells) and most T-cells in atherosclerotic plaques are of the activated CD44^+^ phenotype ^2,62,63^. Sharing similarities with the MESA study, another study found a positive correlation between blood T_memory/effector_ cells and cIMT, and elevated blood T_memory/effector_ cells in patients with chronic stable angina (CSA) or acute myocardial infarction (AMI) ^64^. However, these studies do not take into account the balance between T-cell activation, apoptosis, or senescence. Indeed, T-cell *Abca1/Abcg1* deficiency decreased the expression of pro-inflammatory cytokines in the aorta despite T-cells being more activated. Because it has been reported that uptake of apoptotic T-cells by macrophages may suppress pro-inflammatory gene expression ^57–60^, we reasoned that the enhanced T-cell apoptosis in middle-aged *Ldlr*^-/-^ mice with T-cell *Abca1/Abcg1* deficiency is an anti-inflammatory process ^65^, and may counter-regulate the pro-atherogenic effects of T_memory/effector_ cells.

T-cells in atherosclerotic plaques were increased by 2-fold in middle-aged compared to young *Ldlr*^-/-^ mice, suggesting a more prominent role of T-cells in atherosclerosis in middle-aged mice. Consistent with this hypothesis, previous studies have found different roles of T-cells at different stages of lesion development. Complete CD4^+^ or CD8^+^ T-cell ablation ^66–69^ reduces atherosclerosis, suggesting that T-cells in plaques mainly have a pro-atherogenic role. In early atherosclerotic lesions, CD4^+^ and CD8^+^ T-cells induce macrophage inflammation and monopoiesis mediated by IFNγ ^67^,^69^, while in advanced lesions, CD8^+^ T-cells induce macrophage apoptosis and necrotic core formation mediated by TNFα, granzyme B and perforin ^68^. T-cell *Abca1/Abcg1* deficiency did not affect blood myeloid cells or necrotic core formation, but increased macrophage content in middle-aged *Ldlr*^-/-^ mice, suggestive of less advanced lesions. This was accompanied by decreased inflammation in the aorta. We suggest that the decreased inflammation may be the consequence of uptake of apoptotic T-cells by myeloid cells in the aorta, which is an anti-inflammatory process ^65^.

T-cell *Abca1/Abcg1* deficiency also decreased CD25^+^Foxp3^+^ T_regs_ in para-aortic LNs, which would be suggestive of a pro-atherogenic role of T-cell *Abca1/Abcg1* deficiency. However, *Foxp3* mRNA expression was not affected in aortic T-cells, suggesting no differences in T_regs_ in atherosclerotic plaques. Our *in vitro* studies suggested that the decrease in CD25^+^Foxp3^+^ T_regs_ was entirely due to T-cell apoptosis, an anti-inflammatory process ^65^.

One striking observation was that unlike in young mice, middle-aged *Abca1/Abcg1*-deficient T-cells almost completely lost their ability to proliferate upon TCR stimulation. While this could be the consequence of increased T-cell apoptosis, the senescence marker *p21* was also increased, suggestive of premature T-cell aging. Similar to T-cell *Abca1/Abcg1* deficiency, aging increases T-cell plasma membrane cholesterol accumulation ^16^ and increases T-cell activation, while decreasing peripheral T-cell numbers ^17,18,54^. During aging, thymic involution decreases output of naïve T-cells ^24^. However, T-cells undergo homeostatic proliferation in the periphery giving rise to new T-cells independent of thymic output ^19,70^. We found that T-cells from aged wild-type mice showed increased cholesterol accumulation compared to young wild-type mice, and increased apoptosis upon TCR stimulation, similar to our findings in mice with T-cell *Abca1/Abcg1* deficiency. Together, these data indicate broader physiological relevance of our findings on apoptosis in T-cell *Abca1/Abcg1* deficiency, and suggest that T-cell plasma membrane cholesterol accumulation may account for increased T-cell apoptosis during aging, resulting in a decrease in peripheral T-cell numbers.

Our finding that T-cell *Abca1/Abcg1* deficiency only affected atherosclerotic lesion size in middle-aged *Ldlr*^-/-^ mice is suggestive of a specific effect of *Abca1/Abcg1* on T-cells during aging that affects atherogenesis. T-cell *Abca1/Abcg1* deficiency had similar effects on most T-cell subsets in young and middle-aged mice, but induced T-cell senescence only in middle-aged mice. In humans, T-cell aging due to repeated TCR stimulation is characterized by loss of CD28 ^71^. Human T-cells deficient in CD28 are senescent but still may have an effector function ^72^, and produce high levels of TNFα and IFNγ ^73^. While initial studies reported that CD4^+^CD28^null^ T-cells correlate with unstable angina and acute coronary syndromes in humans ^73–76^, this was recently challenged by a multi-center study with a larger number of patients ^77^. Opposite to the previous findings ^73–76^, in this study ^77^, levels of CD4^+^CD28^null^ T-cells in blood were associated with a lower risk for first time coronary events in a population-based cohort, and did not correlate with cIMT. Based on these findings ^77^, it is unlikely that senescent T-cells contributed to lesion size in our study.

In addition to atherosclerotic plaques, peripheral T-cells in blood, LNs, and spleen were decreased in *T-Abc^dko^Ldlr*^-/-^ mice compared to their littermate controls. A previous study has attributed the decrease in peripheral T-cells in mice with T-cell *Abca1/Abcg1* deficiency or T-cell deficiency of the transcription factor the *liver X receptor* (*LXR*)*α* and *β*, that induces *Abca1* and *Abcg1* expression, to increased thymic CD4^+^ and CD8^+^ T-cell apoptosis ^38^. However, in these studies, the *Abca1* and *Abcg1* floxed genes and also the *Lxrα/Lxrβ* floxed genes were expressed under control of the *LckCre* promoter ^38^. In T-cells, *Lxrβ* is more highly expressed than *Lxrα* ^28^. *CD4CreLxrβ^fl/fl^* mice showed decreased peripheral T-cells with only thymic CD4^+^ T-cells and CD4^+^ T_regs_ being decreased, while thymic CD8^+^ T-cells were unchanged ^29^. Perhaps the *LckCre* promoter resulted in a more complete deficiency of *Abca1* and *Abcg1* in single positive thymic CD4^+^ and CD8^+^ T-cells than the *CD4Cre* promoter that we and others used ^29^, explaining the more pronounced effects on thymic T-cells. The decrease in thymic CD4^+^ and CD8^+^ T-cells in *LckCreLxrα^fl/fl^Lxrβ^fl/fl^* and *LckCreAbca1^fl/fl^Abcg1^fl/fl^* mice was attributed to increased surface expression of the cell death receptor FAS ^38^, which is localized in lipid rafts ^50^. Also, upon TCR stimulation, peripheral CD4^+^ T-cells from *CD4CreLxrβ^fl/fl^* mice showed increased FAS expression and apoptosis; however it was suggested that additional mechanisms may contribute to apoptosis in *CD4CreLxrβ^fl/fl^* T-cells ^29^. Our data in *CD4CreAbca1^fl/fl^Abcg1^fl/fl^Ldlr*^-/-^ mice show that FAS expression was not affected upon TCR stimulation, but Bcl2 expression was decreased. We thus attributed the increased apoptosis in CD4^+^ and CD8^+^ T-cells (the latter mainly in the CD8^+^ T_memory/effector_ T-cells) from *CD4CreAbca1^fl/fl^Abcg1^fl/fl^Ldlr*^-/-^ mice to a decrease in the anti-apoptotic protein Bcl2. This mechanism may also have contributed to the increased apoptosis in CD4^+^ T-cells from *CD4CreLxrβ^fl/fl^* mice.

Similar to *CD4CreAbca1^fl/fl^Abcg1^fl/fl^Ldlr*^-/-^ mice, *CD4CreLxrβ^fl/fl^* mice showed increased spontaneous T-cell activation ^29^. This was attributed to a functional impairment of *CD4CreLxrβ^fl/fl^* T_regs_, and recapitulated in *Foxp3CreLxrβ^fl/fl^* mice, indicating that it was a T_reg_ cell-intrinsic effect ^29^. In these specific *Foxp3CreLxrβ^fl/fl^* T-cells, unlike in other CD4^+^ T-cells, *Abca1* and *Abcg1* expression were not affected by *Lxrβ* deficiency, suggesting that other LXR target genes contributed to the dysfunctional T_reg_ differentiation ^29^. Although peripheral T_regs_ were also decreased in *CD4CreAbca1^fl/fl^Abcg1^fl/fl^Ldlr*^-/-^ mice, there was no specific decrease in the percentage of CD25^+^Foxp3^+^ T-cells, as in *CD4CreLxrβ^fl/fl^* mice ^29^, and the decreases in peripheral T_regs_ were mainly the consequence of an overall increase in T-cell apoptosis. Given that TCR signaling was increased in *CD4CreAbca1^fl/fl^Abcg1^fl/fl^Ldhr*^-/-^ mice, and enhanced TCR signaling promotes T-cell activation ^47^, we postulate that this mechanism accounted for the increase in T-cell activation in *CD4CreAbca1^fl/fl^Abcg1^fl/fl^Ldlr*^-/-^ mice compared to controls.

Several studies have shown that increased plasma membrane cholesterol accumulation enhances TCR signaling ^15^,^28^,^31^. Conversely, impaired cholesterol synthesis in the context of SCAP deficiency suppresses TCR signaling ^46^. *Acat1* deficiency increases plasma membrane cholesterol accumulation and TCR nanoclustering, which enhances TCR signaling ^15^. Cholesterol accumulation also increased TCR nanoclustering in a study employing artificial membranes ^78^; however a later study from the same group showed that binding of cholesterol to TCRβ inhibited TCR signaling ^79^, suggesting rather an inhibitory than a stimulating effect. Upon TCR signaling, the plasma membrane cholesterol content increases ^28,29^, presumably due to suppression of *Abca1* and *Abcg1* expression and increased cholesterol synthesis ^28^. Indeed, our studies have shown that T-cell *Abca1/Abcg1* deficiency increased plasma membrane cholesterol accumulation leading to plasma membrane stiffening and enhanced TCR signaling.

In conclusion, our studies indicate a link between T-cell plasma membrane cholesterol accumulation, T-cell plasma membrane stiffening, TCR signaling, T-cell activation, and T-cell apoptosis in T-cell *Abca1/Abcg1* deficiency, which suppresses aortic inflammation and atherosclerosis in middle-aged, but not young mice. Our studies indicate that plasma membrane cholesterol accumulation not only induces apoptosis, but also a premature T-cell aging phenotype characterized by an increase in T-cell senescence. This suggests that upregulation of *Abca1* and *Abcg1* in T-cells, for example by an LXR agonist, may suppress T-cell apoptosis and senescence, which could be crucial in maintaining T-cell numbers and peripheral T-cell tolerance, especially during aging. Further, statins could also suppress T-cell apoptosis and senescence by suppressing cholesterol synthesis in T-cells. It would be of interest to investigate this further in a large population cohort.

## METHODS

### Experimental mouse models

*Abca1^fl/fl^Abcg1^fl/fl^* (stock 021067), *CD4Cre* (stock 017336), *Ldlr*^-/-^ (stock 002207), and wild-type (stock 000664) mice were purchased from Jackson Laboratories (Bar Harbor, ME). All mice were in the C57Bl6/J background. *Abca1^fl/fl^Abcg1^fl/fl^* (control; *Ldlr^+/+^*) and *CD4Cre* mice were intercrossed to obtain *Abca1^fl/fl^Abcg1^fl/fl^, Abca1^fl/fl^, Abcg1^fl/fl^, CD4CreAbca1^fl/fl^, CD4CreAbcg1^fl/fl^*, and *CD4CreAbca1^fl/fl^Abcg1^fl/fl^* (*T-Abc^dko^*) littermates. *CD4CreAbca1^fl/fl^, CD4CreAbcg1^fl/fl^*, and *CD4CreAbca1^fl/fl^Abcg1^fl/fl^* mice were intercrossed with *Ldlr*^-/-^ mice to generate *Abca1^fl/fl^Ldlr^-/-^, Abcg1^fl/fl^Ldlr^-/-^, Abca1^fl/fl^Abcg1^fl/fl^Ldlr*^-/-^ (all *Ldlr*^-/-^), *CD4CreAbca1^fl/fl^Ldlr*^-/-^ (*T-Abca1^ko^Ldlr*^-/-^), *CD4CreAbcg1^fl/fl^Ldlr*^-/-^ (*T-Abcg1^ko^Ldlr*^-/-^), and *CD4CreAbca1^fl/fl^Abcg1^fl/fl^Ldlr*^-/-^ (*T-Abc^dko^Ldlr*^-/-^) littermates. Mice were fed a chow diet (V1554-703, Ssniff Spezialdiäten GmbH) or a Western-type diet (WTD; 40% fat, 0.15% cholesterol; D12079B, Research Diets).

For all studies, littermates were used, and mice were housed under standard laboratory conditions with a light cycle of 12 hours and ad libitum water and food (chow diet or WTD). Water and cages were autoclaved. Cages were changed once weekly, and the health status of the mice was monitored based on body weight, coat, and behavior. The mouse genotype did not cause changes in body weight (mouse body weight between 20-40 g, depending on gender and age) and health. Female littermates (atherosclerosis studies) or littermates from both sexes were randomly assigned to experimental groups, unless stated otherwise. The number of the mice used for the experiments are indicated for each experiment in the figure legends. In general, n=3-18 mice were used per group and experiments were repeated at least once if n=3 mice were used per experiment to confirm the reproducibility of the results. No inclusion or exclusion criteria were used. All protocols were approved by the Institutional Animal Care and Use Committee from the University of Groningen (Groningen, the Netherlands).

### T-cell and thymocyte isolations

Spleens were isolated from control, *T-Abc^dko^*, *Ldlr*^-/-^, *T-Abc^dko^Ldlr*^-/-^, and young (3 months) and aged (24 months) wild-type mice, mashed on a 40 μm strainer, and red blood cells (RBCs) were lysed (Lysing buffer, BD Bioscience). Para-aortic lymph nodes (LNs) were isolated from *Ldlr*^-/-^ and *T-Abc^dko^Ldlr*^-/-^ mice and mashed on a 40 μm strainer in PBS. Splenic and LN homogenates were then centrifuged, washed, and resuspended in MACS buffer (PBS containing 0.5% BSA and 2mM EDTA). Dead cells were removed using the Dead Cell Removal kit (Miltenyi Biotec), and total T-cells were isolated using the Pan T cell kit (Miltenyi Biotec). CD4^+^ and CD8^+^ T-cells were isolated using CD4 and CD8 coated magnetic beads (Miltenyi Biotec), respectively. For thymocyte isolations, *Ldlr*^-/-^ and *T-Abc^dko^Ldlr*^-/-^ thymi were isolated and mashed on a 40 μm strainer in PBS.

### Flow cytometry and total white blood cell counts

Blood samples were collected from control, *T-Abc^dko^, Ldlr^-/-^, T-Abca1^sko^Ldlr^-/-^, T-Abcg1^sko^Ldlr^-/-^, T-Abc^dko^Ldlr^-/-^*, young and aged wild-type mice by tail bleeding into Eppendorf tubes containing EDTA. Total white blood cell counts were assessed using the Medonic CD620 hematology analyzer (Boule Medical). For flow cytometry, tubes were kept at 4°C for the whole procedure unless stated otherwise. RBCs were lysed (Lysing buffer, BD Bioscience), and white blood cells were centrifuged, washed, and resuspended in FACS buffer (HBSS containing 0.1% BSA and 0.5mM EDTA). For analysis of blood T-cells, cells were stained with a cocktail of antibodies against TCRβ-PB (Biolegend), CD8-FITC, and CD4-APC (eBioscience). CD4^+^ T-cells were identified as TCRβ^+^CD4^+^ and CD8^+^ T-cells as TCRβ^+^CD8^+^. For analysis of naïve and activated blood T-cells, cells were stained with a cocktail of antibodies against TCRβ-PB (Biolegend), CD8-FITC, CD44-PE-Cy7, and CD62L-APC (eBioscience). Among these populations, T_naive_ cells were identified as CD44^-^CD62L^+^, T_central memory_ cells as CD8^+^CD44^+^CD62L^+^, and T_memory/effector_ cells as CD44^+^CD62L^-^. The same stainings were carried out on para-aortic LN and splenic homogenates. For analysis of thymic T-cells (for *Ldlr*^-/-^ and *T-Abc^dko^Ldlr*^-/-^ mice), thymic homogenates were stained with CD4-APC and CD8-FITC (eBioscience). For analysis of blood myeloid cells (for *Ldlr*^-/-^ and *T-Abc^dko^Ldlr*^-/-^ mice), cells were stained with a cocktail of antibodies against CD45-PE-Cy7, Ly6-C/G-PerCP-Cy5.5 (BD Biosciences), and CD115-PE (Biolegend). Monocytes were identified as CD45^+^CD115^+^ and further separated into Ly6-C^hi^ and Ly6-C^lo^ subsets, and neutrophils were identified as CD45^+^CD115^-^Ly6-C/G^hi^.

For analyses of T-cell subsets and T-cell exhaustion, para-aortic LN and thymic homogenates from *Ldlr*^-/-^ and *T-Abc^dko^Ldlr*^-/-^ mice were used. For analysis of T_regulatory_ cells, LN and thymic homogenates were stained with a cocktail of antibodies against CD4-FITC and CD25-APC (for LN T_regulatory_ cells; Mouse Regulatory T Cell Staining Kit #1, eBioscience) or CD4-PB (Biolegend), CD8-FITC, and CD25-PECy7 (eBioscience) (for thymic T_regulatory_ cells). Cells were then fixed and permeabilized using the Foxp3/Transcription Factor Staining Buffer Set (eBioscience), and subsequently stained with Foxp3-PE (for LN T_regulatory_ cells) or Foxp3-APC (for thymic T_regulatory_ cells) or their isotype control (eBioscience). T_regulatory_ cells were identified as CD8^-^ CD4^+^CD25^+^Foxp3^+^ and CD8^-^CD4^+^CD25^-^Foxp3^+^. For analysis of naïve and activated T_regulatory_ cells, LN homogenates were stained with a cocktail of antibodies against CD4-PE (Biolegend), CD25-PECy7, CD44-PB, and CD62L-FITC (eBioscience). Cells were then fixed and permeabilized using the Foxp3/Transcription Factor Staining Buffer Set (eBioscience), and subsequently stained with Foxp3-APC or its isotype control (eBioscience). Naïve T_regulatory_ cells were identified as CD4^+^CD25^+^Foxp3^+^CD44^-^CD62L^+^ and CD4^+^CD25^-^Foxp3^+^CD44^-^CD62L^+^. Activated T_regulatory_ cells were identified as CD4^+^CD25^+^Foxp3^+^CD44^+^CD62L^-^ and CD4^+^CD25^-^ Foxp3^+^CD44^+^CD62L^-^ (T_memory/effector_). For analysis of T_follicular helper_ cells and total PD1^+^ T-cells, LN homogenates were stained with a cocktail of antibodies against CD4-APC-Cy7 (Biolegend), CD44-PE-Cy7, CD62L-APC, PD1-PB (eBioscience), and CXCR5-FITC (Biolegend). T_follicular helper_ cells were defined as CD4^+^CD44^+^CD62^-^CXCR5^+^PD1^+^ and total PD1^+^ T-cells as CD4^+^PD1^+^.

For analysis of T_bet_^+^ cells, LN homogenates were stained with a cocktail of antibodies against TCRβ-PB (Biolegend), CD4-APC, and CD8-FITC (eBioscience), then fixed and permeabilized, and subsequently stained with T_bet_-PE or its isotype control (eBioscience). T_bet_^+^ cells were identified as TCRβ^+^CD4^+^T_bet_^+^ and TCRβ^+^CD8^+^T_bet_^+^. For analysis of naïve and activated T_bet_^+^ cells, LN homogenates were stained with a cocktail of antibodies against TCRβ-PB (Biolegend), CD8-PE, CD44-PECy7, and CD62L-APC (eBioscience), then fixed and permeabilized, and subsequently stained with T_bet_-AF488 or its isotype control (eBioscience). Naive T_bet_^+^ cells were identified as TCRβ^+^CD4^+^T_bet_^+^CD44^-^CD62L^+^ and TCRβ^+^CD8^+^T_bet_^+^CD44^-^CD62L^+^. Activated T_bet_^+^ cells were identified as TCRβ^+^CD4^+^T_bet_^+^CD44^+^CD62L^-^, TCRβ^+^CD8^+^T_bet_^+^CD44^+^CD62L^-^ (T_memory/effector_) and TCRβ^+^CD8^+^T_bet_^+^CD44^+^CD62L^+^ (T_central memory_) cells. For analysis of the exhaustion marker CTLA4, LN homogenates were stained with a cocktail of antibodies against CD4-APC, CD8-FITC, and CTLA4 (eBioscience). For analysis of the exhaustion markers TIM3 and LAG3, LN homogenates were stained with a cocktail of antibodies against TCRβ-PerCPCy5.5 (Biolegend), CD4-FITC, CD8-PE (eBioscience), and TIM3-APC or LAG3-APC (Biolegend). For analysis of the exhaustion marker Eomes, LN homogenates were stained with a cocktail of antibodies against TCRβ-PB (Biolegend), CD8-APC, and CD4-FITC (eBioscience), then fixed and permeabilized, and subsequently stained with Eomes-PE or its isotype control (eBioscience). CTLA4, TIM3, and LAG3 mean fluorescence intensity (mfi) on CD4^+^ and CD8^+^ T-cells, and Eomes mfi in CD8^+^ T-cells was assessed by flow cytometry.

All samples were analyzed on LSRII (BD Biosciences), running FACSDiVa software (BD Biosciences). The data were analyzed using the FlowJo software (FlowJo).

### Filipin staining

For analysis of free cholesterol in T-cells, blood leukocytes (from *Ldlr^-/-^, T-Abc^dko^Ldlr^-/-^*, young and aged wild-type mice) and thymocytes (from *Ldlr*^-/-^ and *T-Abc^dko^Ldlr^-/^* mice) were used. Blood leukocytes were stained with a cocktail of antibodies against TCRβ-APC (Biolegend), CD8-PE, and CD4-FITC (eBioscience) and thymocytes with a cocktail of antibodies against CD4-APC and CD8-PE (eBioscience). Cells were then centrifuged, washed, and stained with 50 μg/mL Filipin III (Sigma-Aldrich) in 10% FCS/PBS for 45 min at room temperature and then washed twice with PBS. Samples were analyzed by flow cytometry immediately after staining. Filipin III mfi on blood TCRβ^+^CD4^+^ and TCRβ^+^CD8^+^ cells, and on thymic CD4^+^CD8^+^, CD4^+^, and CD8^+^ T-cells was assessed by flow cytometry.

### Choleratoxin B staining

For analysis of T-cell lipid rafts, blood leukocytes from *Ldlr*^-/-^ and *T-Abc^dko^Ldlr*^-/-^ mice were stained with a cocktail of antibodies against TCRβ-PB (Biolegend), CD8-PE, and CD4-APC (eBioscience). Cells were then centrifuged, washed, and stained with 200 ng/mL choleratoxin B-FITC (Sigma-Aldrich) for 1 h at room temperature. Samples were analyzed by flow cytometry immediately after staining. Choleratoxin B mfi on TCRβ^+^CD4^+^ and TCRβ^+^CD8^+^ cells was assessed by flow cytometry, and is an indirect indicator of lipid rafts.

### *In vitro* T-cell apoptosis assays

Splenic CD4^+^ and CD8^+^ T-cells from *Ldlr*^-/-^ and *T-Abc^dko^Ldlr*^-/-^ mice were isolated and labeled using Incucyte Caspase3/7 Red Reagent (Sartorius). Cells were cultured in RPMI 1640 medium (Gibco) supplemented with 10% FCS, 1% pen-strep, and 50 μM β-mercaptoethanol at 37°C, 5% CO_2_. To induce apoptosis, cells were stimulated with 5 μg/mL plate-bound αCD3 (Biolegend) and 100 U/mL IL-2 (Peprotech) for 12 h. For Incucyte experiments, images were taken every 2 h. For image analysis, background signal was removed using the Incucyte ZOOM software (Sartorius) and the number of total cells and Caspase3^+^/7^+^ cells per image were quantified using Image J software (NIH) in a blinded fashion, *i.e*. the observer was unaware of the genotypes. Alternatively, splenic T-cells from *Ldlr^-/-^, T-Abc^dko^Ldlr^-/-^*, young wild-type, and aged wild-type mice were isolated and labeled using Incucyte Caspase3/7 Red Reagent (Sartorius). Cells were stimulated with 5 μg/mL plate-bound αCD3 (Biolegend) and 100 U/mL IL-2 (Peprotech) for 12 h to induce apoptosis. Cells were then harvested, stained with CD4-PB (Biolegend) or CD8-PB (eBioscience), and analyzed by flow cytometry. Caspase3/7 mfi in CD4^+^ and CD8^+^ T-cells was assessed by flow cytometry.

### FAS and Bcl2 assays

Splenic T-cells from *Ldlr*^-/-^ and *T-Abc^dko^Ldlr*^-/-^ mice were isolated as described above and stimulated with 5 μg/mL plate-bound αCD3 (Biolegend) and 100 U/mL IL-2 (Peprotech) for 12 h to induce apoptosis. Cells were harvested and stained with a cocktail of antibodies against CD4-PB (Biolegend), CD8-PE, and FAS-AF488 (for FAS analysis; eBioscience) or TCRβ-PB (Biolegend), CD8-PE, CD44-PE-Cy7, and CD62L-APC (for Bcl2 analysis; eBioscience). For Bcl2 analysis, cells were then fixed and permeabilized, and subsequently stained with Bcl2-FITC or its isotype control (eBioscience). Samples were analyzed by flow cytometry. FAS mfi on CD4^+^ and CD8^+^ T-cells was assessed. Percentages of CD4^+^Bcl2^+^, CD4^+^Bcl2^-^, CD8^+^ CD44^+^CD62L^-^Bcl2^+^, and CD8^+^CD44^+^CD62L^-^Bcl2^-^ T-cells were quantified. Apoptosis was confirmed by flow cytometry using the caspase 3/7 assay as described above.

### Mitochondrial ROS assay

Splenic T-cells from *Ldlr*^-/-^ and *T-Abc^dko^Ldlr*^-/-^ mice were isolated as described above and stimulated with 5 μg/mL plate-bound αCD3 (Biolegend) and 100 U/mL IL-2 (Peprotech) for 12 h. Cells were harvested and stained with 2 μM MitoSOX Red Mitochondrial Superoxide Indicator or 50 nM MitoTracker Green FM (Invitrogen; carbocyanine-based dye) for 30 min at 37°C. For MitoSOX (eBioscience) analysis, cells were stained with a cocktail of antibodies against CD44-PE-Cy7, CD62L-APC, CD4-PB (Biolegend) or CD8-PB. For MitoTracker (eBioscience) analysis, cells were stained with a cocktail of antibodies against CD44-PB, CD62L-APC, CD4-PE (Biolegend) or CD8-PE. MitoSOX and Mitotracker mfi on CD4^+^ and CD8^+^ T_naive_, CD4^+^ and CD8^+^ T_memory/effector_, and CD8+ T_central memory_ cells was assessed by flow cytometry. To correct the levels of mitochondrial ROS for mitochondrial mass, the MitoSOX/Mitotracker ratio was calculated.

### *In vitro* T-cell proliferation assay

Splenic T-cells from *Ldlr^-/-^, T-Abc^dko^Ldlr^-/-^*, young wild-type, and aged wild-type mice were isolated and labeled with 10 μM CFSE-PB (Invitrogen) and stimulated with Mouse T-Activator CD3/CD28 Dynabeads (ratio beads to cells 1:2) in RPMI 1640 medium supplemented with 10% FCS, 1% pen-strep, and 50 μM β-mercaptoethanol. Cells were incubated at 37°C for 72 h. Three days later, beads were removed, T-cells were stained using CD4 and CD8 antibodies as described above, and CFSE-PB dilution was assessed by flow cytometry. Using the Proliferation Modeling tool in the FlowJo software (FlowJo), histograms of CFSE-PB dilution in CD4^+^ and CD8^+^ T-cells were obtained where each cell division was depicted as a separate peak. The number of CD4^+^ and CD8^+^ T-cells in each division was automatically calculated by the software after designing gates for each peak. To calculate the percentage of CD4^+^ and CD8^+^ T-cells in each division, the number of CD4^+^ and CD8^+^ T-cells in one division was divided by the total number of CD4^+^ and CD8^+^ T-cells in all divisions, respectively.

### *In vitro* T_regulatory cell_ differentiation assay

Splenic T-cells from *Ldlr*^-/-^ and *T-Abc^dko^Ldlr*^-/-^ mice were isolated and stimulated with Mouse T-Activator CD3/CD28 Dynabeads (ratio beads to cells 1:2) in the presence of 100 U/mL IL-2 (Peprotech) and 5 ng/mL TGFβ1 (Peprotech) in RPMI 1640 medium supplemented with 10% FCS, 1% pen-strep, and 50 μM β-mercaptoethanol. Cells were incubated at 37°C for 72 h. Three days later, beads were removed, T-cells were stained using CD4, CD25, and Foxp3 antibodies as described above, and samples were analyzed by flow cytometry. T_regulatory_ cells were identified as CD4^+^CD25^+^Foxp3^+^.

### Aortic cell isolations

Aortas were isolated from *Ldlr*^-/-^ and *T-Abc^dko^Ldlr*^-/-^ mice and digested for 1 hour at 37°C using an enzyme mixture that contained liberase TH (4 U/mL), DNAse I (40 U/mL), and hyaluronidase (60 U/mL). Aortic homogenates were then centrifuged, washed, and resuspended in MACS buffer. Total aortic CD3^+^ T-cells and CD3^-^ cells were isolated using the Pan T cell kit (Miltenyi Biotec).

### RNA extraction from splenic and aortic cells

Splenic total T-cells, CD4^+^, and CD8^+^ T-cells from *Abca1^fl/fl^Abcg1^fl/fl^, CD4CreAbca1^fl/fl^Abcg1^fl/fl^, Ldlr*^-/-^, *T-Abc^dko^Ldlr*^-/-^, young wild-type, and aged wild-type mice and aortic total CD3^+^ T-cells and CD3^-^ cells from *Ldlr*^-/-^ and *T-Abc^dko^Ldlr*^-/-^ mice were isolated as described above and resuspended directly into RLT buffer. RNA was then extracted using the RNeasy Mini Kit (Qiagen) and cDNA was synthesized using the Transcriptor Universal cDNA Master kit (Roche). *Abca1, Abcg1* (in splenic total T-cells), *p21* (in splenic CD4^+^ and CD8^+^ T-cells and aortic total CD3^+^ T-cells), *Foxp3, Tgfβ, Bcl2*, (in aortic total CD3^+^ T-cells), *Il-10, Tnfa* (in aortic total CD3^+^ T-cells and CD3^-^ cells), *Cd11b, Mcp1, Il-6*, and *Il-1β* (in aortic CD3^-^ cells) mRNA levels were assessed by qPCR using QuantStudio 7 Flex Real-Time PCR System (Applied Biosystems), and initial differences in RNA quantity were corrected for using the housekeeping genes *36b4* and *Cyclophilin B*.

### Plasma total cholesterol

Blood samples were collected from *Ldlr*^-/-^ and *T-Abc^dko^Ldlr*^-/-^ mice by tail bleeding. Plasma was separated by centrifugation. To assess lipoprotein cholesterol distribution by fast performance liquid chromatography (FPLC), pooled plasma (n=14-15 per pool) was injected onto a Superose 6 HR 10/300 GL column (GE Healthcare) and eluted at a constant flow rate of 500 μL/min. Fractions were assayed for cholesterol using an enzymatic kit from Roche enzymatic kit and the Cholesterol standard FS (DiaSys Diagnostic Systems) for the calibration curve, which were also used to measure total plasma cholesterol.

### Atherosclerotic lesion analysis

Female *Ldlr*^-/-^ and *T-Abċ^dko^Ldlr*^-/-^ mice were fed a WTD for 10 weeks or a chow diet for 28 weeks or 12-13 months. After the indicated period of time on diet, mice were sacrificed and the hearts were isolated and fixed in phosphate-buffered formalin. Hearts were dehydrated and embedded in paraffin, and cross-sectioned throughout the aortic root area (4 μm sections). Haematoxylin-eosin (H&E) staining was performed on the sections and the average from 6 sections (40 μm distance between sections) for each animal was used to determine lesion size. Lesion size was quantified by morphometric analysis using the ImageJ software (NIH). Necrotic core area was determined as acellular area, lacking nuclei and cytoplasm, under the fibrous cap of lesions from H&E stained sections. Necrotic core area was differentiated from regions of dense fibrous scars by the presence of macrophage debris. To assess the number of T-cells in atherosclerotic lesions, sections were stained with a CD3 antibody (Dako) and counterstained with haematoxylin. T-cells were identified as cells with brown plasma membrane CD3 staining and the hematoxylin staining still visible. Quantification of the number of T-cells per section was performed in a blinded fashion, *i.e*. the observer was unaware of the genotypes. To assess the macrophage content of atherosclerotic lesions, sections were stained with a Mac-2 primary antibody (Cedarlane) and an anti-rat IgG-AF488 secondary antibody (Invitrogen). Sections were mounted using ProLong Gold Antifade Mountant with DAPI (Thermofisher;) and imaged using an AxioObserver Z1 compound microscope (10x objective; AxioCam MRm3 CCD camera; Carl Zeiss) running Zen software (Zeiss). The Mac-2 positive area was quantified using the ImageJ software (NIH) and expressed as % of total lesion area.

### Oil Red O staining

*Ldlr-/-* and *T-Abc^dko^Ldlr*^-/-^ mice were sacrificed and para-aortic LNs were isolated and embedded into OCT compound on dry ice. Frozen cross-sections were made and stained with Oil Red O (Sigma-Aldrich) to assess neutral lipid accumulation. Sections were counterstained with haematoxylin. Quantification of the number of lipid droplets was performed in a blinded fashion, *i.e*. the observer was unaware of the genotypes.

### Gas Chromatography – Mass Spectrometry

Splenic T-cells from *Ldlr*^-/-^ and *T-Abc^dko^Ldlr*^-/-^ mice were isolated as described above and pelleted. Cholestanol-D5 was added to the samples as internal standard and cholesterol was extracted using hexane. Total and free cholesterol content was determined by Gas Chromatography - Mass Spectrometry (7890B GS system, 5973 MS system, and 7693A automatic liquid sampler from Agilent; positive chemical ionization mode with 5% ammonia in methane as reaction gas). A polar DB-WAXetr (30m x 0.25 mm x 0.25 μm) column was used. Cholesteryl ester content was calculated by subtracting free cholesterol from total cholesterol. Cholesterol content was normalized to cellular protein content. Cellular protein content was measured by the Lowry assay.

### T-cell plasma membrane stiffness by fluorescence-lifetime imaging microscopy (FLIM)

Splenic CD4^+^ and CD8^+^ T-cells from *Ldlr*^-/-^ and *T-Abc^dko^Ldlr*^-/-^ mice were isolated as described above. Cells were stained with BODIPY C10 (kind gift from Dr. Ulf Diederichsen, Georg-August-Universität Göttingen, Germany; 4 μM; 30 min; on ice). Cells were then washed twice with phenol red free RPMI 1640 medium and seeded in Poly-L-Lysine (PLL) coated μ-Slide 8 Well Glass Bottom Chambers (Ibidi). Cells were incubated at 37°C for 30 min to attach to the glass, and then BODIPY C10 fluorescence lifetime was assessed by FLIM microscopy. All images were collected with a MicroTime 200 microscope (PicoQuant) equipped with an Olympus (100x/1,4) oil immersion objective. Images were acquired using the SymPhoTime 64 software (PicoQuant). Data analysis of the FLIM images was performed using the open-source FLIMfit software ^80^. Further image loading options included 2×2 spatial binning and time bins 1563 (32ps/bin). After loading FLIM images, the software’s default settings were used. BODIPY C10 fluorescence lifetime of a whole cell (with LDs) or of the plasma membrane without including lipid droplets (without LDs) was measured using the tool “Fit Selected Decay”. Per mouse, 43-76 CD4^+^ and CD8^+^ T-cells were analyzed.

### Confocal microscopy

Splenic CD4^+^ and CD8^+^ T-cells from *Ldlr*^-/-^ and *T-Abc^dko^Ldlr*^-/-^ mice were isolated as described above. Cells were seeded on PLL-coated glass coverslips and incubated at 37°C for 1 h to attach to the glass.

#### Lipid droplet staining

For imaging of lipid droplets in CD4^+^ and CD8^+^ T-cells, cells were fixed (4% PFA; 15 min; 4°C), permeabilized (CLSM buffer = PBS containing 3% BSA, 10 mM glycine, 0.1% saponin; 30 min; room temperature), and stained with BODIPY 493/503 (Invitrogen; 7.5 μM in CLSM buffer; 30 min; room temperature). After two PBS washes, cells were mounted using ProLong Gold Antifade Mountant with DAPI (Vector Laboratories).

#### Lipid droplet and mitochondrial staining

To assess lipid droplets in mitochondria, CD4^+^ and CD8^+^ T-cells were stained with MitoTracker Red CMXRos (Invitrogen; derivative of X-rosamine; 200 nM in serum free medium; 30 min; 37°C), fixed (3.7% PFA; 15 min; 37°C), permeabilized (CLSM buffer; 30 min; room temperature), and stained with BODIPY 493/503 (Invitrogen; 7.5 μM in CLSM buffer; 30 min; room temperature). After two PBS washes, cells were mounted using ProLong Gold Antifade Mountant with DAPI (Vector Laboratories).

#### Free cholesterol and mitochondrial staining

To assess free cholesterol in mitochondria, CD4^+^ and CD8^+^ T-cells were stained with MitoTracker Red CMXRos (Invitrogen; 200 nM in serum free medium; 30 min; 37°C), fixed (3% PFA; 1 h; room temperature), and stained with filipin (50 μg/mL in 10% FCS in PBS; 2 h; room temperature). After three PBS washes, cells were mounted using ProLong Gold Antifade Mountant (Vector Laboratories).

#### Free cholesterol and lysosomal staining

To assess free cholesterol in lysosomes, CD4^+^ and CD8^+^ T-cells were stained with LysoTracker Red DND-99 (Invitrogen; 5 μM; 1 h; 37°C), fixed (3% PFA; 1 h; room temperature), and stained with filipin (50 μg/mL in 10% FCS in PBS; 2 h; room temperature). After three PBS washes, cells were mounted using ProLong Gold Antifade Mountant (Vector Laboratories).

All samples were imaged on the Zeiss LSM800 microscope with immersion oil lens (63X,1.4 N.A) running Zen software (Zeiss). Background signal was removed using Zen software and images were analyzed using Image J software (NIH).

### Transmission electron microscopy

Splenic CD4^+^ and CD8^+^ T-cells from *Ldlr*^-/-^ and *T-Abc^dko^Ldlr*^-/-^ mice were isolated as described above. Cells were seeded on PLL-coated glass coverslips placed in 24-well plates and incubated at 37°C for 1 h to attach to the glass. Cells were then fixed with 2% glutaraldehyde (Sigma-Aldrich) in PB (0.1 M phosphate buffer, pH 7.4) for 60 min at room temperature. After three PB washes, cells were post-fixed with 1% osmium tetroxide in PB for 60 min at room temperature. Cells were incubated overnight in 0.5% uranyl acetate, dehydrated with graded steps of ethanol (50%, 70%, 96%, 100%), and embedded in Epon resin. Sections of 70 nm thickness were made and stained with 2% uranyl acetate solution and lead citrate solution. Stained sections were then examined using a CM12 transmission electron microscope (Phillips).

### Quantification and statistical analysis

All data are presented as means ± SEM. In each experiment, n defines the number of mice included. The statistical parameters (n, mean, SEM) can be found within the figure legends. Two-tailed Student’s t-test was used to define differences between two datasets. To define differences between three or four datasets, One-way Analysis of Variance (ANOVA) was used with a Bonferroni multiple comparison post-test. The criterion for significance was set at *p*<0.05. Statistical analyses were performed using GraphPad Prism 9 (San Diego, California).

## DATA AVAILABILITY

The authors declare that the data supporting the findings of this study are available within the paper and its supplementary information files.

## ACKNOWLEDGMENTS

M.W. is supported by VIDI grant 917.15.350 from the Netherlands Organization of Scientific Research (NWO), and a Rosalind Franklin Fellowship from the University of Groningen.

## AUTHORS CONTRIBUTIONS

V.B.: Conceptualization, Methodology, Investigation, Writing – Original Draft preparation. A.M.L.R., R.d.B., E.G., A.F-S., A.G.G., A.M-G., M.H.K., N.J.K., A.P., M.L-M., A.d.B.: Investigation. S.M., F.B.: Resources, Investigation. B.v.d.S, A.K., L.Y-C., G.v.d.B.: Conceptualization, Writing – Review & Editing. M.W.: Conceptualization, Methodology, Investigation, Writing – Review & Editing, Supervision, Funding Acquisition.

## DECLARATION OF INTERESTS

The authors declare no competing interests.

## ADDITIONAL INFORMATION

Correspondence and requests for materials should be addressed to M.W.

## Notes

### Competing Interest Statement

The authors have declared no competing interest.

