## Supplemental Figures for "T-cell Abca1 and Abcg1 cholesterol efflux pathways suppress T-cell apoptosis and senescence and increase atherosclerosis in middle-aged *Ldlr*^-/-^ mice"

Supplementary Fig. 1

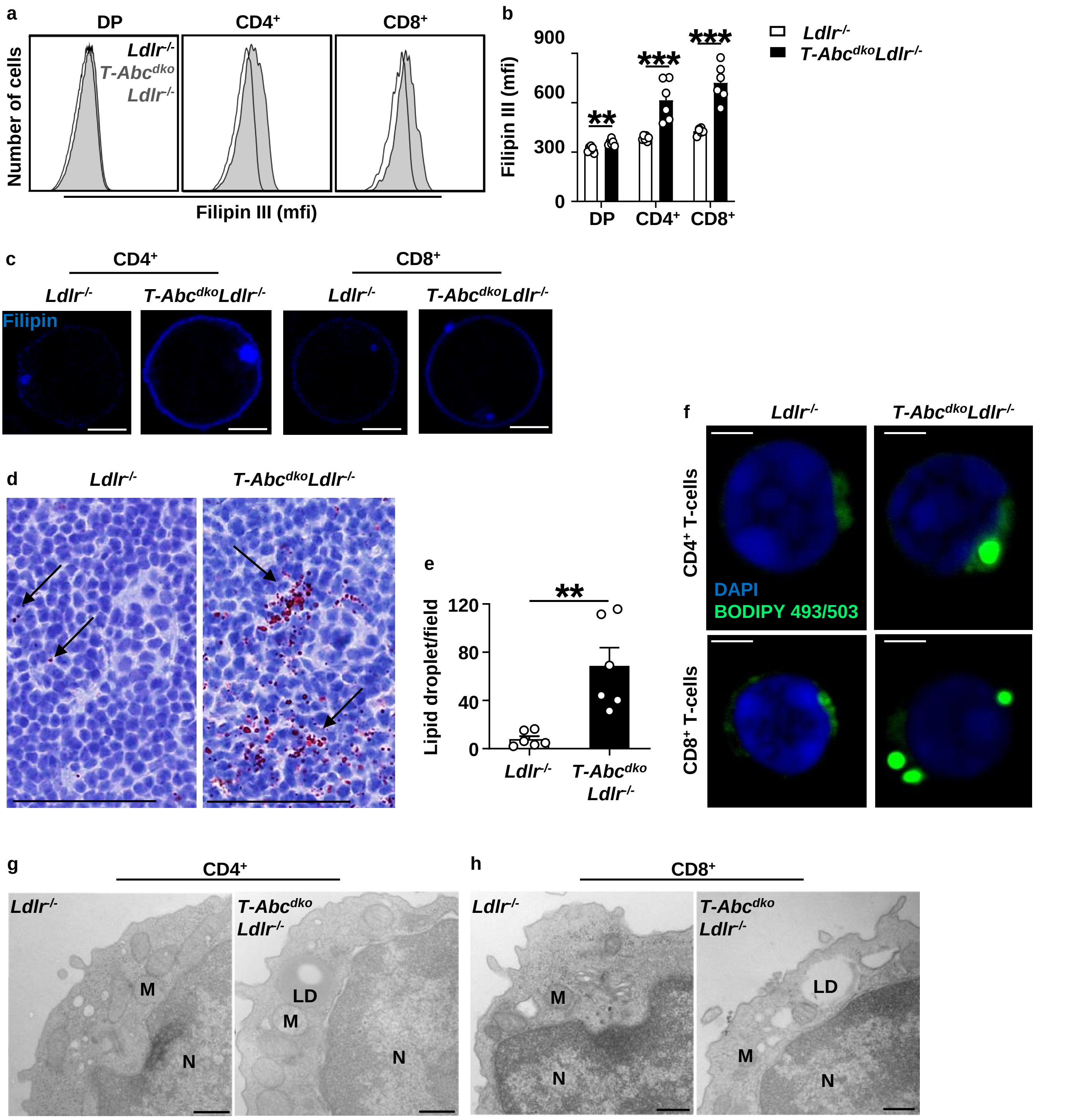


**Supplementary Fig. 1 T-cell *Abca1/Abcg1* deficiency increases cholesterol accumulation.** *T-Abc*$^{dko}$*Ldlr*$^{-/-}$ and *Ldlr*$^{-/-}$ mice were fed a chow diet. Thymi, spleens, and para-aortic lymph nodes (LNs) were collected. (**a-b**) Filipin staining on thymic T-cells was assessed by flow cytometry. (**a**) Representative flow cytometry plots and (**b**) quantification. n=6. (**c**) Splenic CD4$^+$ and CD8$^+$ T-cells were isolated, fixed, stained with filipin, and analyzed by confocal microscopy. Representative pictures are shown. Scale bar represents 2 µm. (**d-e**) Para-aortic LNs were embedded in OCT and frozen sections were stained for Oil Red O. Representative pictures are shown (**d**). Lipid droplets are depicted by arrows (**d**) and quantified (**e**). n=6. Scale bar represents 60 µm. For panels **b** and **e**, error bars represent standard error of the mean (SEM). n indicates biological replicates. *p* value was determined by unpaired two-tailed Student's t-test. ***p*<0.01, ****p*<0.001. (**f**) Splenic CD4$^+$ and CD8$^+$ T-cells were fixed, permeabilized, and stained with BODIPY 493/503 and DAPI. Samples were analyzed by confocal microscopy and representative pictures are shown. Scale bar represents 2 µm. (**g-h**) Representative transmission electron microscopy images for CD4$^+$ (**g**) and CD8$^+$ (**h**) T-cells are shown. LD = lipid droplet. M = mitochondria. N = nucleus. Scale bar represents 500 nm.

**Supplementary Fig. 2**

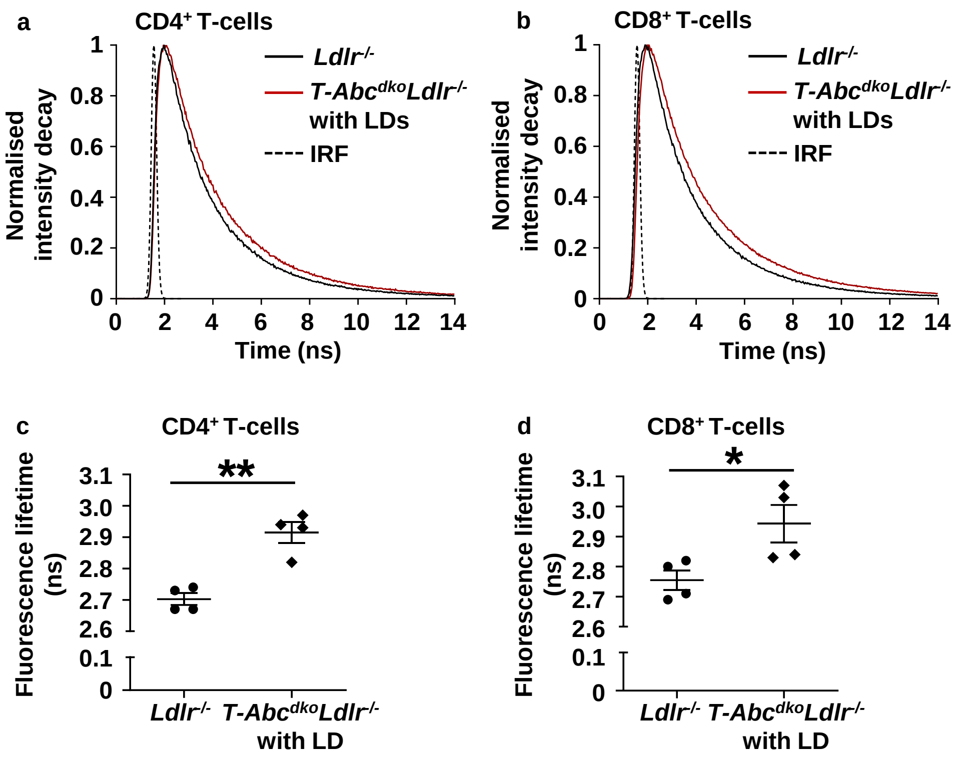


**Supplementary Fig. 2 T-cell *Abca1/Abcg1* deficiency increases cell stiffness.** *T-Abc*$^{dko}$*Ldlr*$^{-/-}$ and *Ldlr*$^{-/-}$ mice were fed a chow diet. Spleens were collected and CD4$^{+}$ and CD8$^{+}$ T-cells were isolated. CD4$^{+}$ and CD8$^{+}$ T-cells were stained with BODIPY C10 and analyzed by Fluorescence-Lifetime Imaging Microscopy (FLIM). Fluorescence lifetime decay curves for CD4$^{+}$ (**a**) and CD8$^{+}$ (**b**) T-cells are shown and quantified in the plasma membrane and lipid droplet (with LDs) (**c-d**). Each data point represents an individual mouse. LD = lipid droplet. (**a-b**) Dashed lines: fits with mono-exponential decay functions convoluted with the instrument response function (IRF). n=4. For all panels, error bars represent SEM. n indicates biological replicates. *p* value was determined by unpaired two-tailed Student's t-test. **p*<0.05, ***p*<0.01.

Supplementary Fig. 3

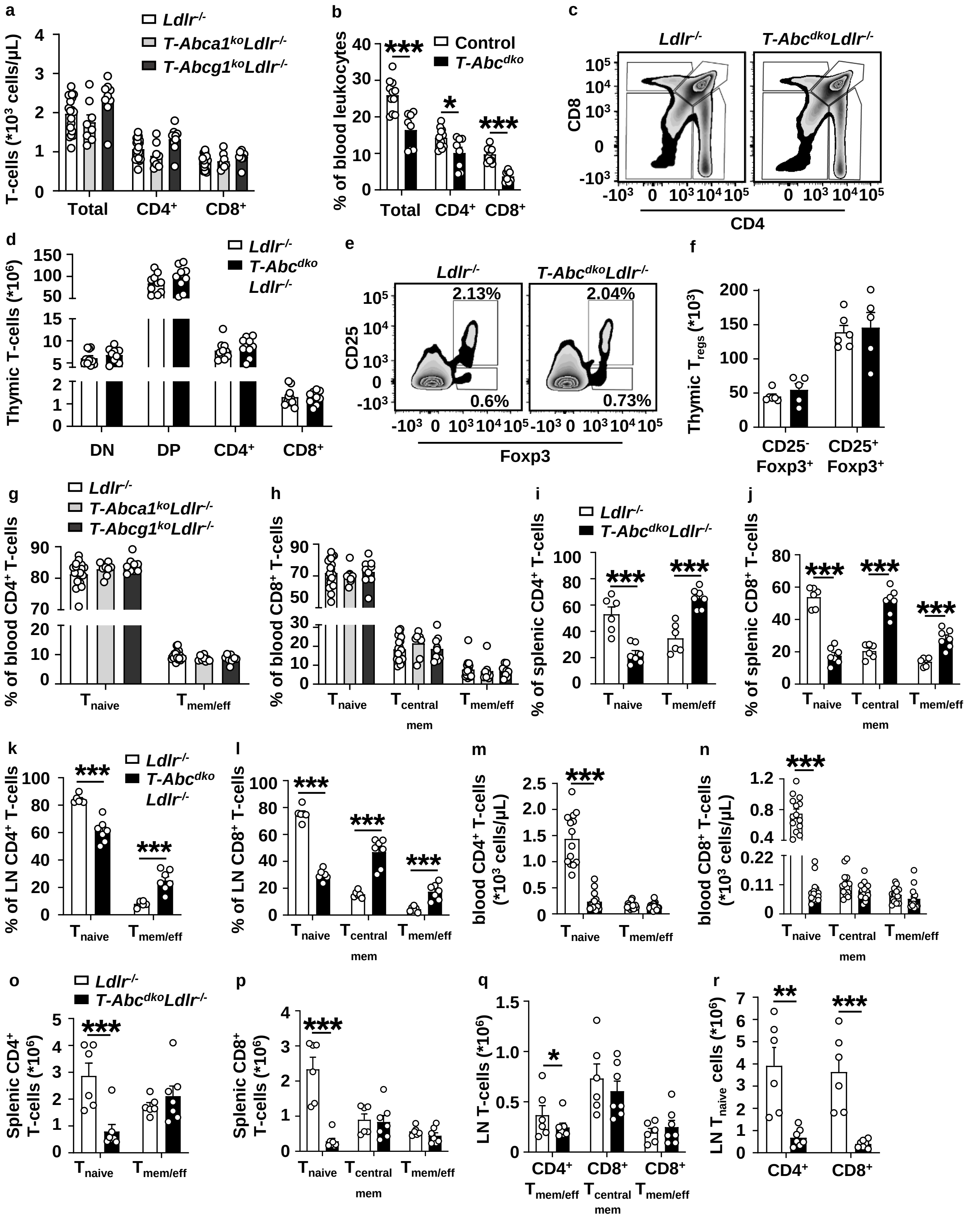

**Supplementary Fig. 3 T-cell *Abca1/Abcg1* deficiency does not affect thymic T-cells and increases T-cell activation.** Control**,** *T-Abc$^{dko}$, Ldlr$^{-/-}$*, *T-Abca1$^{sko}$Ldlr$^{-/-}$*, *T-Abcg1$^{sko}$Ldlr$^{-/-}$*, and *T-Abc$^{dko}$Ldlr$^{-/-}$* mice were fed a chow diet. Blood, thymi, spleens, and para-aortic LNs were collected. Cells were stained with the indicated antibodies and analyzed by flow cytometry. (**a**) TCRβ$^{+}$ (total), CD4$^{+}$, and CD8$^{+}$ T-cells in blood from *Ldlr$^{-/-}$*, *T-Abca1$^{sko}$ Ldlr$^{-/-}$*, and *T-Abcg1$^{sko}$ Ldlr$^{-/-}$* mice. n=8-19. (**b**) TCRβ$^{+}$ (total), CD4$^{+}$, and CD8$^{+}$ T-cells in blood from control (*Ldlr$^{+/+}$*) and *T-Abc$^{dko}$* mice. n=8-11. (**c-d**) Representative flow cytometry plots of thymic CD4$^{-}$CD8$^{-}$ (DN), CD4$^{+}$CD8$^{+}$ (DP), CD4$^{+}$, and CD8$^{+}$ T-cells are shown (**c**), corrected for total thymocytes, and quantified (**d**). n=9-10. (**e-f**) Representative flow cytometry plots of thymic CD25$^{-}$Foxp3$^{+}$ and CD25$^{+}$Foxp3$^{+}$ $T_{regs}$ are shown (**e**), corrected for total thymocytes, and quantified (**f**). n=6. (**g-h**) CD4$^{+}$ CD44$^{-}$CD62L$^{+}$ ($T_{naive}$) and CD44$^{+}$CD62L$^{-}$ ($T_{mem/eff}$) (**g**), and CD8$^{+}$ $T_{naive}$, $T_{mem/eff}$, and CD8$^{+}$CD44$^{+}$CD62L$^{+}$ ($T_{central\ mem}$) (**h**) cells in blood from *Ldlr$^{-/-}$*, *T-Abca1$^{sko}$ Ldlr$^{-/-}$*, and *T-Abcg1$^{sko}$ Ldlr$^{-/-}$* mice. n=8-19. (**i-l**) Percentages of splenic (**i-j**) and para-aortic LN (**k-l**) CD4$^{+}$ $T_{naive}$ and $T_{mem/eff}$ cells (**i, k**) and CD8$^{+}$ $T_{naive}$, $T_{mem/eff}$, and $T_{central\ mem}$ cells (**j, l**). n=6-7. (**m-p**) Numbers of blood (**m-n**) and splenic (**o-p**) CD4$^{+}$ $T_{naive}$ and $T_{mem/eff}$ cells (**m, o**) and CD8$^{+}$ $T_{naive}$, $T_{mem/eff}$, and $T_{central\ mem}$ cells (**n, p**). n=6-7. (**q-r**) Numbers of para-aortic LN CD4$^{+}$ and CD8$^{+}$ $T_{mem/eff}$, and CD8$^{+}$ $T_{central\ mem}$ cells (**q**) and CD4$^{+}$ and CD8$^{+}$ $T_{naive}$ cells (**r**). n=6-7. For all panels, error bars represent SEM. n indicates biological replicates. *p* value was determined by unpaired two-tailed Student's t-test (**b, d, f, i-r**) or one-way ANOVA with Bonferroni post-test (**a, g-h**) . **p*<0.05, ***p*<0.01, ****p*<0.001.

Supplementary Fig. 4

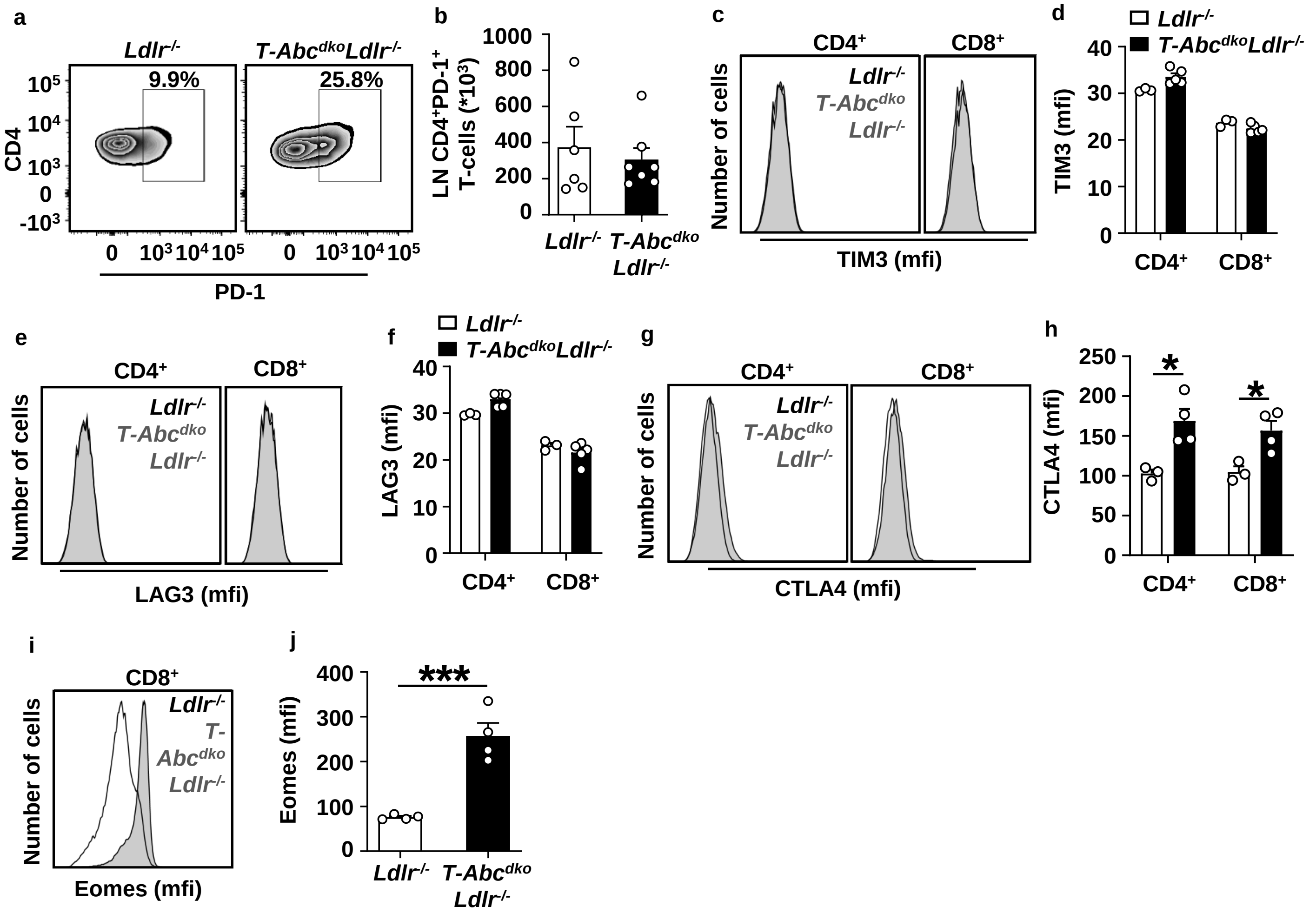


**Supplementary Fig. 4 Effects of T-cell *Abca1/Abcg1* deficiency on T-cell exhaustion.** *T-Abc*$^{dko}$*Ldlr*$^{-/-}$ and *Ldlr*$^{-/-}$ mice were fed a chow diet. Para-aortic LNs were isolated and cells were stained with the indicated antibodies and analyzed by flow cytometry. Representative flow cytometry plots of CD4$^{+}$PD-1$^{+}$ T-cells (**a**), and TIM3 (**c**), LAG3 (**e**), and CTLA4 (**g**) in CD4$^{+}$ and CD8$^{+}$ T-cells and Eomes (**I**) in CD8$^{+}$ T-cells are shown and quantified (**b, d, f, h, j**). n=3-7. For all panels, error bars represent SEM. n indicates biological replicates. *p* value was determined by unpaired two-tailed Student's t-test. *$p<0.05$, ***$p<0.001$.

**Supplementary Fig. 5**

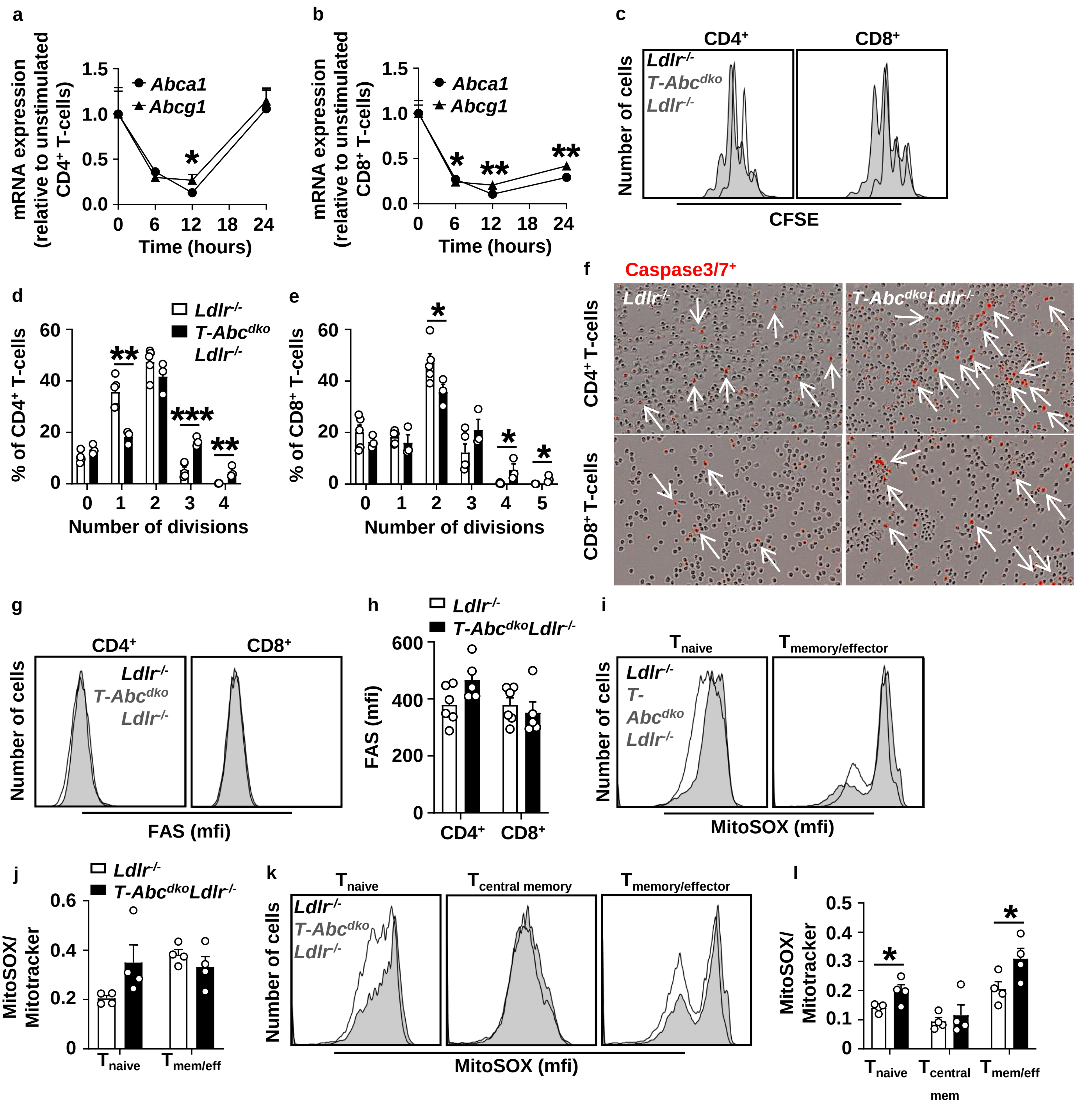


**Supplementary Fig. 5 Effects of T-cell *Abca1/Abcg1* deficiency on responses downstream of the T-cell receptor.** *T-Abc$^{dko}$Ldlr$^{-/-}$* and *Ldlr$^{-/-}$* mice were fed a chow diet. Spleens were collected and CD4$^{+}$ and CD8$^{+}$ T-cells were isolated. (**a-b**) CD4$^{+}$ and CD8$^{+}$ T-cells from *Ldlr$^{-/-}$* mice were stimulated with αCD3/αCD28 beads for the indicated time. RNA was extracted and *Abca1* and *Abcg1* mRNA expression were measured in CD4$^{+}$ (**a**) and CD8$^{+}$ (**b**) T-cells by qPCR. n=4-5. (**c-e**) T-cells were labeled with CFSE and stimulated with αCD3/αCD28 beads. CFSE dilution was measured at 72 h after stimulation by flow cytometry. Representative CFSE dilutions are shown (**c**). The number of divisions were quantified for CD4$^{+}$ (**d**) and CD8$^{+}$ T-cells (**e**). n=3-5. (**f-l**) CD4$^{+}$ and CD8$^{+}$ T-cells were isolated and stimulated with αCD3 and IL-2 for 12 h (**f-l**) with concomitant staining for cleaved Caspase3/7 (**f**). (**f**) CD4$^{+}$ and CD8$^{+}$ T-cells acquiring Caspase 3$^{+}$/7$^{+}$ staining over time were assessed using the Incucyte system. Snapshots at 10 h after stimulation are shown. Caspase 3$^{+}$/7$^{+}$ CD4$^{+}$ and CD8$^{+}$ T-cells are depicted by arrows. (**g-h**) Representative flow cytometry plots of FAS expression on CD4$^{+}$ and CD8$^{+}$ T-cells are shown (**g**) and quantified (**h**). n=5-6. (**i-j**) Representative flow cytometry plots of MitoSOX levels in CD4$^{+}$ $T_{naive}$ and $T_{mem/eff}$ cells are shown (**i**) and quantified (**j**). n=4. (**k-l**) Representative flow cytometry plots of MitoSOX levels in CD8$^{+}$ $T_{naive}$, $T_{mem/eff}$, and $T_{central\ mem}$ cells are shown (**k**) and quantified (**l**). n=4. MitoSOX values in (**j**) and (**l**) have been corrected for MitoTracker Green levels. For all panels, error bars represent SEM. n indicates biological replicates. *p* value was determined by unpaired two-tailed Student's t-test. *$p<0.05$, **$p<0.01$, ***$p<0.001$.

**Supplementary Fig. 6**

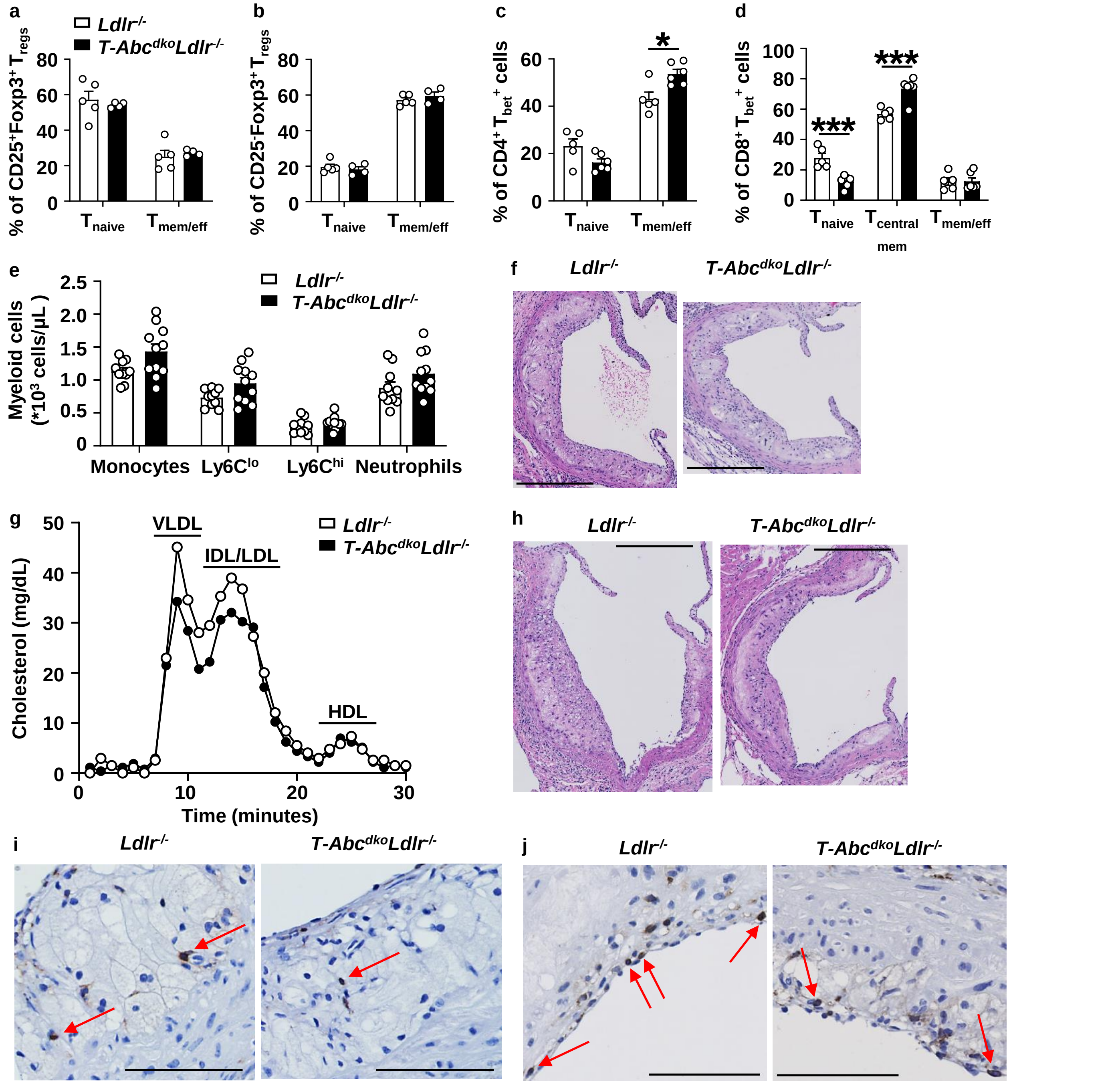


**Supplementary Fig. 6 T-cell *Abca1/Abcg1* deficiency does not affect atherosclerotic lesion size but decreases T-cells in atherosclerotic plaques of *Ldlr*$^{-/-}$ mice.** Female *T-Abc*$^{dko}$*Ldlr*$^{-/-}$ and *Ldlr*$^{-/-}$ mice were fed a chow diet for 28 weeks (**a-d, h, j**) or a Western-type diet (WTD) for 10 weeks (**e-g, i**). (**a-d**) Para-aortic LNs were isolated and cells were stained with the indicated antibodies and analyzed by flow cytometry. (**a-b**) Quantification of $T_{naive}$ and $T_{mem/eff}$ CD25$^{+}$Foxp3$^{+}$ (**a**) and CD25$^{-}$Foxp3$^{+}$ (**b**) $T_{regs}$. n=4-5. (**c-d**) Quantification of $T_{naive}$ and $T_{mem/eff}$ CD4$^{+}$T$_{bet}^{+}$ (**c**) and $T_{naive}$, $T_{mem/eff}$, and $T_{central\ mem}$ CD8$^{+}$T$_{bet}^{+}$ (**d**) cells. n=5-6. (**e**) Blood was collected and white blood cells were stained with the indicated antibodies and analyzed by flow cytometry. Numbers of total monocytes, Ly6C$^{lo}$ and Ly6C$^{hi}$ monocyte subsets, and neutrophils. n=10-11. For panels **a-e**, error bars represent SEM. n indicates biological replicates. *p* value was determined by unpaired two-tailed Student's t-test. *$p$<0.05, ***$p$<0.001. (**f, h, i-j**) Hearts were isolated and sections of the aortic root were prepared, and stained for hematoxylin-eosin (H&E) (**f, h**) or CD3 (**i-j**). (**f, h**) Representative pictures of H&E staining on the aortic root from WTD-fed (**f**) and chow diet-fed (**h**) mice. Scale bar represents 300 µm. (**g**) Lipoprotein fractions from pooled plasma samples (n=14-15 per pool) of mice fed WTD were separated using fast performance liquid chromatography and cholesterol levels were determined. HDL = high-density-lipoprotein. IDL = intermediate-density-lipoprotein. LDL = low-density-lipoprotein. VLDL = very low-density-lipoprotein. (**i, j**) Representative pictures of CD3 staining on atherosclerotic lesions from WTD-fed (**i**) and chow diet-fed (**j**) mice. T-cells were identified as cells with brown plasma membrane CD3 staining and the hematoxylin staining still visible. CD3$^{+}$ cells are depicted by arrows. Scale bar represents 80 µm.

Supplementary Fig. 7

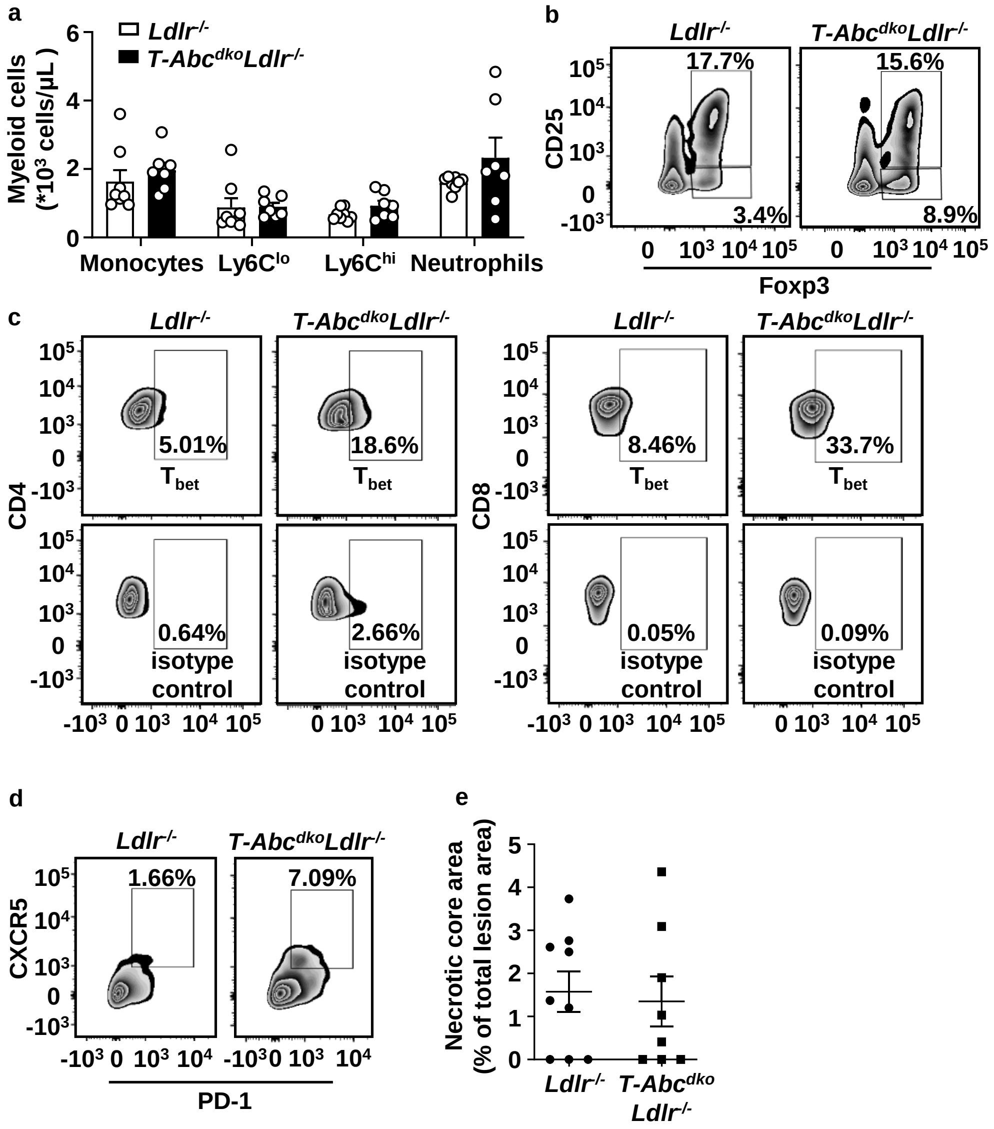


**Supplementary Fig. 7 Effects of T-cell *Abca1/Abcg1* deficiency on blood myeloid cells, LN T-cell subsets, and necrotic core area in atherosclerotic plaques of middle-aged *Ldlr*$^{-/-}$ mice.** *T-Abc*$^{dko}$*Ldlr*$^{-/-}$ and *Ldlr*$^{-/-}$ mice were fed a chow diet for 12-13 months. (**a**) Blood was collected and white blood cells were stained with the indicated antibodies and analyzed by flow cytometry. Numbers of total monocytes, Ly6C$^{lo}$ and Ly6C$^{hi}$ monocyte subsets, and neutrophils. n=7-8. (**b-d**) Para-aortic LNs were isolated and cells were stained with the indicated antibodies and analyzed by flow cytometry. (**b**) Representative flow cytometry plots of CD25$^{-}$Foxp3$^{+}$ and CD25$^{+}$Foxp3$^{+}$ T$_{regs}$. (**c**) Representative flow cytometry plots of CD4$^{+}$ and CD8$^{+}$T$_{bet}^{+}$ T-cells and their respective isotype control. (**d**) Representative flow cytometry plots of CD4$^{+}$CD44$^{+}$CD62L$^{-}$CXCR5$^{+}$PD1$^{+}$ T$_{FH}$ cells. (**e**) Hearts were isolated. Sections of the aortic root were prepared and stained for H&E. Necrotic core area corrected for total atherosclerotic lesion area. n=8-11. Each data point represents an individual mouse. For all panels, error bars represent SEM. n indicates biological replicates.

**Supplementary Table 1. Plasma cholesterol levels in *T-Abc*$^{dko}$*Ldlr*$^{-/-}$ and *Ldlr*$^{-/-}$ mice fed Western-type diet (WTD) or chow diet**

| Genotype | Diet | Plasma cholesterol levels (mg/dL) |
|---|---|---|
| *Ldlr*$^{-/-}$ | WTD | 919.9 ± 52.4 |
| *T-Abc*$^{dko}$*Ldlr*$^{-/-}$ | WTD | 780.2* ± 32 |
| *Ldlr*$^{-/-}$ | Chow diet (28 weeks) | 283.8 ± 15.1 |
| *T-Abc*$^{dko}$*Ldlr*$^{-/-}$ | Chow diet (28 weeks) | 295.3 ± 16.2 |
| *Ldlr*$^{-/-}$ | Chow diet (12-13 months) | 255.8 ± 14.8 |
| *T-Abc*$^{dko}$*Ldlr*$^{-/-}$ | Chow diet (12-13 months) | 249.1 ± 18.6 |

Plasma cholesterol levels were determined using an enzymatic kit. n=15 on WTD. n=16-18 on chow diet for 28 weeks. n=10-11 on chow diet for 12-13 months. For all mouse groups, ± SEM is shown. n indicates biological replicates. *p* value was determined by unpaired two-tailed Student’s t-test. *$p<0.05$.

**Supplementary table 2. Reagents and resources.**

| REAGENT or RESOURCE | SOURCE | IDENTIFIER |
|---|---|---|
| Antibodies | | |
| Rat anti-mouse CD115-PE | Biolegend | Cat# 135506; RRID:AB_1937253; clone AFS98 |
| Rat anti-mouse Ly6-C/G-PerCP-Cy5.5 | BD Biosciences | Cat# 561103; RRID:AB_10562568; clone RB6-8C5 |
| Rat anti-mouse CD45-APC-Cy7 | BD Biosciences | Cat# 557659; RRID:AB_396774; clone 30-F11 |
| Rat anti-mouse CD25-PECy7 | eBioscience | Cat# 25-0251-82; RRID:AB_469608; clone PC61.5 |
| Rat anti-mouse CD4-PB | Biolegend | Cat# 100427; RRID:AB_493646; clone GK1.5 |
| Rat anti-mouse CD4-APC | eBioscience | Cat# 17-0041-82; RRID:AB_469320; clone GK1.5 |
| Rat anti-mouse CD4-APC-Cy7 | Biolegend | Cat# 100413; RRID:AB_312698; clone GK1.5 |
| Rat anti-mouse CD4-PE | Biolegend | Cat# 100407; RRID:AB_312692; clone GK1.5 |
| Rat anti-mouse CD4-FITC | eBioscience | Cat# 11-0042-82; RRID:AB_464896; clone RM4-5 |
| Rat anti-mouse CD8-PB (eFluor 450) | eBioscience | Cat# 48-0083-82; RRID:AB_11218504; clone eBioH35-17.2 (H35-17.2) |
| Rat anti-mouse CD8-FITC | eBioscience | Cat# 11-0083-82; RRID:AB_657764; clone eBioH35-17.2 (H35-17.2) |
| Rat anti-mouse CD8-PE | eBioscience | Cat# 12-0083-82; RRID:AB_657767; clone eBioH35-17.2 (H35-17.2) |
| Armenian hamster anti-mouse TCRβ-PB | Biolegend | Cat# 109226; RRID:AB_1027649; clone H57-597 |
| Armenian hamster anti-mouse TCRβ-APC | Biolegend | Cat# 109211; RRID:AB_313434; clone H57-597 |
| Armenian hamster anti-mouse TCRβ-PerCP-Cy5.5 | Biolegend | Cat# 109227; RRID:AB_1575176; clone H57-597 |

| | | |
|---|---|---|
| Armenian hamster anti-mouse PD1-PB (eFluor 450) | eBioscience | Cat# 48-9985; RRID:AB_2574138; clone J43 |
| Rat anti-mouse PD1-PE | Biolegend | Cat# 109103; RRID:AB_313420; clone RMP1-30 |
| Mouse anti mouse $T_{bet}$-PE | eBioscience | Cat# 12-5825; RRID:AB_925761; clone eBio4B10 (4B10) |
| Mouse IgG1 kappa isotype control-PE ($T_{bet}$- and Eomes-PE) | eBioscience | Cat# 12-4714-42; RRID:AB_470060; clone P3.6.2.8.1 |
| Mouse anti mouse $T_{bet}$-AF488 | eBioscience | Cat# 53-5825-80; RRID:AB_2815215; clone eBio4B10 (4B10) |
| Mouse IgG1 kappa isotype control-AF488 ($T_{bet}$-AF488) | eBioscience | Cat# 53-4714-80; RRID:AB_470230; clone P3.6.2.8.1 |
| Rat anti-mouse Eomes-PE | eBioscience | Cat# 12-4875-82; RRID:AB_1603275; clone Dan11mag |
| Rat anti-mouse CXCR5-FITC | Biolegend | Cat# 145519; RRID:AB_2562865; clone L138D7 |
| Rat anti-mouse CD44-PB (eFluor 450) | eBioscience | Cat# 48-0441-82; RRID:AB_1272246; clone IM7 |
| Rat anti-mouse CD44-PE-Cy7 | eBioscience | Cat# 25-0441-82; RRID:AB_469623; clone IM7 |
| Rat anti-mouse CD62L-APC | eBioscience | Cat# 17-0621-82; RRID:AB_469410; clone MEL-14 |
| Rat anti-mouse CD62L-FITC | Biolegend | Cat# 104405; RRID:AB_313092; clone MEL-14 |
| Rat anti-mouse Foxp3-APC | eBioscience | Cat# 17-5773-80; RRID:AB_469456; clone FJK-16s |
| Rat IgG2a kappa isotype control-APC (Foxp3-APC) | eBioscience | Cat# 17-4321-81; RRID:AB_470181; clone eBR2a |
| Mouse anti-mouse Fas-AF488 | eBioscience | Cat# 53-0951-82; RRID:AB_10671269; clone 15A7 |
| Mouse anti-mouse Bcl2-FITC | eBioscience | Cat# 11-6992-42; RRID:AB_10734060; clone 10C4 |
| Mouse IgG1 kappa isotype control-FITC (Bcl2-FITC) | eBioscience | Cat# 11-4714-82; RRID:AB_470022 |

| | | |
|---|---|---|
| Armenian hamster anti-mouse CTLA4-APC | eBioscience | Cat# 17-1522-82; RRID:AB_2016700; clone UC10-4B9 |
| Rat anti-mouse TIM3-APC | Biolegend | Cat# 134007; RRID:AB_2562997; clone B8.2C12 |
| Rat anti-mouse LAG3-APC | Biolegend | Cat# 125209; RRID:AB_10639935; clone C9B7W |
| Rabbit anti-human CD3 (for histology) | Dako | Cat# A0452; RRID:AB_2335677 |
| Rat Anti-Mouse/Human Mac-2 (Galectin-3) (for histology) | Cedarlane | Cat# CL8942AP; RRID:AB_2534074;clone M3/38 |
| Goat anti-Rat IgG (H+L) Cross-Adsorbed Secondary Antibody, AF488 (for histology) | Invitrogen | Cat# A-11006; RRID:AB_10060357 |
| LEAF Purified rat anti-mouse CD3 (for stimulation) | Biolegend | Cat# 100208; RRID:AB_312665; clone 17A2 |
| Chemicals and Recombinant Proteins | | |
| Incucyte Caspase3/7 Red Reagent | Sartorius | Cat# 4704 |
| Cell Proliferation Dye eFluor 450 (CFSE-PB) | eBioscience | Cat# 65-0842-85 |
| Filipin III | Sigma-Aldrich | Cat# F4767 |
| Filipin complex | Sigma-Aldrich | Cat# F9765 |
| Choleratoxin B-FITC | Sigma-Aldrich | Cat# C1655 |
| MitoSOX Red Mitochondrial Superoxide Indicator | Invitrogen | Cat# M36008 |
| Mitotracker Green FM | Invitrogen | Cat# M7514 |
| MitoTracker Red CMXRos | Invitrogen | Cat# M7512 |
| Bodipy 493/503 | Invitrogen | Cat# D3922 |
| Bodipy C10 | Ulf Diederichsen Lab, Georg-August-Universität Göttingen, Germany; [1] | N/A |
| VECTASHIELD Antifade Mounting Medium with DAPI | Vector Laboratories | Cat# H-1200 |
| VECTASHIELD Antifade Mounting Medium | Vector Laboratories | Cat# H-1000 |
| ProLong Gold Antifade Mountant with DAPI | Invitrogen | Cat# P36935 |
| LysoTracker Red DND-99 | Invitrogen | Cat# L7528 |
| Oil Red O | Sigma-Aldrich | Cat# O0625 |
| Recombinant murine IL-2 | Peprotech | Cat# 212-12 |
| Animal-Free Recombinant Human TGFβ1 (CHO derived) | Peprotech | Cat# AF-100-21C |
| Critical Commercial Assays | | |
| Dynabeads Mouse T-Activator CD3/CD28 for T-Cell Expansion and Activation | Gibco | Cat# 11452D |
| Dead Cell Removal Kit | Miltenyi Biotec | Cat# 130-090-101 |

| | | |
|---|---|---|
| Pan T Cell Isolation Kit II, mouse | Miltenyi Biotec | Cat# 130-095-130 |
| CD4 (L3T4) MicroBeads, mouse | Miltenyi Biotec | Cat# 130-117-043 |
| CD8a (Ly-2) MicroBeads, mouse | Miltenyi Biotec | Cat# 130-117-044 |
| Cholesterol reagent | Roche | Cat# 11489232 |
| Cholesterol Standard FS | Diasys Diagnostic Systems | Cat# 113009910026 |
| Mouse Regulatory T Cell Staining Kit #1 | eBioscience | Cat# 88-8111 |
| Foxp3/Transcription Factor Staining Buffer Set | eBioscience | Cat# 00-5523 |
| Lysing Buffer | BD Biosciences | Cat# 555899 |
| Experimental Models: Organisms/Strains | | |
| Mouse: Abca1$^{fl/fl}$Abcg1$^{fl/fl}$:<br>B6.Cg-Abca1$^{tm1Jp}$ Abcg1$^{tm1Tall}$/J | The Jackson Laboratory | Cat# JAX:021067;<br>RRID:IMSR_JAX:021067 |
| Mouse: CD4Cre:<br>Tg(Cd4-cre)1Cwi/BfluJ | The Jackson Laboratory | Cat# JAX:017336;<br>RRID:IMSR_JAX:017336 |
| Mouse: Ldlr$^{-/-}$:<br>B6.129S7-Ldlr$^{tm1Her}$/J | The Jackson Laboratory | Cat# JAX:002207;<br>RRID:IMSR_JAX:002207 |
| Mouse: wild-type: C57BL/6J | The Jackson Laboratory | Cat# JAX:000664;<br>RRID:IMSR_JAX:000664 |
| Oligonucleotides | | |
| Mouse Abca1fl/fl forward primer<br>5'-GTGAATGGGCAATTCGCAAACT-3' | [2] | N/A |
| Mouse Abca1fl/fl reverse primer<br>5'-AGATCTCCCCTCCTTGACAATGC-3' | [2] | N/A |
| Mouse Abcg1fl/fl forward primer<br>5'-TGTTCAGGAGGCCATGATGGT-3' | [2] | N/A |
| Mouse Abcg1fl/fl reverse primer<br>5'-TGGCCAGGCGTTTCCG-3' T | [2] | N/A |
| Mouse p21 forward primer<br>5'-TTGCCAGCAGAATAAAAGGTG-3' | [3] | N/A |
| Mouse p21 reverse primer<br>5'-TTTGCTCCTGTGCGGAAC-3' | [3] | N/A |
| Mouse Tnfα forward primer<br>5'-GTAGCCCACGTCGTAGCAAAC-3' | [4] | N/A |
| Mouse Tnfα reverse primer<br>5'-AGTTGGTTGTCTTTGAGATCCATG-3' | [4] | N/A |
| Mouse Foxp3 forward primer<br>5'-TCTTGCCAAGCTGGAAGACT-3' | [5] | N/A |
| Mouse Foxp3 reverse primer<br>5'-GGGGTTCAAGGAAGAAGAGG-3' | [5] | N/A |
| Mouse Il-10 forward primer<br>5'-GCTCTTACTGACTGGCATGAG-3' | [6] | N/A |
| Mouse Il-10 reverse primer<br>5'-CGCAGCTCTAGGAGCATGTG-3' | [6] | N/A |
| Mouse Tgfβ forward primer<br>5'-GCCCTTCCTGCTCCTCATG-3' | [7] | N/A |
| Mouse Tgfβ reverse primer<br>5'-CCGCACACAGCAGTTCTTCTC-3' | [7] | N/A |

| | | |
|---|---|---|
| Mouse Bcl2 forward primer 5'-GAACTGGGGGAGGATTGTGG-3' | [8] | N/A |
| Mouse Bcl2 reverse primer 5'-ACCTACCCAGCCTCCGTTAT-3' | [8] | N/A |
| Mouse Cd11b forward primer 5'-TCAGAGAATGTCCTCAGCAG-3' | [4] | N/A |
| Mouse CD11b reverse primer 5'-TGAGACAAACTCCTTCATCTTC-3' | [4] | N/A |
| Mouse Mcp1 forward primer 5'-GCTGGAGAGCTACAAGAGGATCA-3' | [7] | N/A |
| Mouse Mcp1 reverse primer 5'-ACAGACCTCTCTCTTGAGCTTGGT-3' | [7] | N/A |
| Mouse Il-6 forward primer 5'-CTGCAAGAGACTTCCATCCAGTT-3' | [9] | N/A |
| Mouse Il-6 reverse primer 5'-AGGGAAGGCCGTGGTTGT-3' | [9] | N/A |
| Mouse Il-1β forward primer 5'-TGCAGCTGGAGAGTGTGG-3' | [4] | N/A |
| Mouse Il-1β reverse primer 5'-TGCTTGTGAGGTGCTGATG-3' | [4] | N/A |
| Software and Algorithms | | |
| FACS DiVa software V8.0.3 | BD Biosciences | https://www.bdbiosciences.com/us/instruments/research/software/ flow-cytometry-acquisition/bdfacsdiva-software/m/111112/overview |
| FlowJo version 10.6.2 | FlowJo | https://www.flowjo.com/solutions/flowjo/downloads |
| GraphPad Prism version 9 | GraphPad Software | https://www.graphpad.com/scientificsoftware/prism/ |
| ImageJ | NIH | https://imagej.nih.gov/ij/index.html |
| IncuCyte ZOOM 2018A | Sartorius | https://www.essenbioscience.com/en/resources/incucyte-zoom-resources-support/software-modules-incucyte-zoom/ |
| SymPhoTime 64 version 2.6 | PicoQuant | https://www.picoquant.com/products/category/software/symphotime-64-fluorescence-lifetime-imaging-and-correlation-software |
| FLIMfit 5.1.1 | FLIMfit, Open Microscopy Environment | http://flimfit.org/downloads/ |

| | | |
|---|---|---|
| Zen | Zeiss | https://www.zeiss.com/microscopy/us/products/microscope-software/zen-lite.html |
